# Navigating the salinity gradient: Individual variation in habitat use and migration of European eel revealed by otolith microchemistry

**DOI:** 10.64898/2026.06.24.734179

**Authors:** Philip Jacobson, Lisa Spotowitz, Yvette Heimbrand, Elin Myrenås, Rob van Gemert, Josefin Sundin

## Abstract

Knowledge regarding variation in habitat use among individuals is crucial for understanding population dynamics and for management and conservation measures. This is especially important for diadromous fishes that shift between habitats, being affected by external pressures and environmental change in different habitats over ontogeny. Herer, we assessed individual variation in habitat use of European eel along a salinity gradient, ranging from fully marine to freshwater in northern Europe, using otolith microchemistry data from >3600 eel together with established time-series segmentation and clustering methods. We show that eel display high degree of individual variation in habitat use. Assigned life-histories included coastal resident, freshwater resident, and coastal and freshwater habitat shifting individuals. Coastal resident eels were observed in a large range of salinities. Given the widespread occurrence of migration barriers in freshwater, it is unknown whether the coastal resident eel preferred that habitat, or if it was the only available habitat for them. Our findings nonetheless highlight the need to include coastal habitats when assessing population development and silver eel production of the critically endangered European eel.

## Introduction

Knowledge regarding habitat use of fish is fundamental for understanding how they will be affected by environmental change. Many fish species are characterized by complex life-histories as they often undergo ontogenetic shifts in both resource and habitat use with a high degree of variation both within and between species and populations (Werner & Gilliam, 1984; Klementsen et al. 2003; Dodson et al., 2013; Klemetsen, 2013). Accounting for intraspecific variation in habitat use is important for efficient management and conservation measures (e.g. McDowall 1999). This is because individual variation in habitat use can buffer habitat-specific changes since not all individuals will be affected equally by changes in external pressures (Werner & Gilliam, 1984; Miller & Rudolf, 2011), which in turn, can have profound effects on population and community dynamics (Persson & DeRoos, 2003; Knight et al., 2005; Schreiber & Rudolf, 2008). Acquiring such knowledge is however challenging, in particular for fishes with ontogenetic habitat shifts characterized by long-distance migrations in association with large distribution areas e.g., Atlantic bluefin tuna, *Thynnus thynnus* (Horton et al., 2020), Atlantic salmon, S*almo salar* (Rikardsen et al., 2021) and European eel, *Anguilla anguilla* (Moriarty & Dekker 1997; Righton et al., 2016).

Diadromous fishes are particularly important and difficult to study in this regard since they are generally dependent on marine/brackish and freshwater systems to complete their life cycle (Gross et al., 1988; Klementsen et al., 2003). The shift between habitats is not always straightforward and may not simply consist of one migration episode between habitats to, for example, complete spawning. Rather, great variation in habitat use over ontogeny, both between and within populations, have been documented for many diadromous fish species (e.g. Quinn & Meyers, 2004; Daverat et al., 2006; Moore et al., 2014; Jacobson et al., 2020; Rikardsen et al., 2021). Furthermore, many diadromous fish populations show declining numbers and even local extinction due to the combined negative effects of e.g. decreased connectivity caused by migration barriers, overfishing, hydropower turbine mortality, climate change, pollution, aquaculture, urbanization, changes in land-use, and invasive species (e.g. Limburg & Waldman, 2009; Thorstad et al. 2021; Gillson et al. 2022; Waldman & Quinn, 2022; Dadswell et al. 2022; Dempson et al. 2024; Ohlberger et al. 2025).

The European eel (*Anguilla anguilla* L.) is a critically endangered, semelparous, panmictic fish species with a complex life cycle and a large distribution area including the Sargasso Sea, Iceland, northern Norway and Russia to southern Europe and the northern parts of Africa, the Azores, the Baltic-, Mediterranean– and the Black Sea (Moriarty & Dekker, 1997; Palm et al., 2009; Pike et al., 2020; Enbody et al., 2021). The leptocephalus larvae hatch in the Sargasso Sea from where they are transported via oceanic currents towards the eastern North Atlantic continental shelf (Schmidt 1912; 1922; Miller et al., 2019). Upon arrival at the continental shelf, the leptocephalus larvae metamorphose into transparent glass eel, colonizing coastal and freshwater habitats where they enter the growth life stage known as yellow eel (Tesch, 2003). The duration of the yellow eel life stage is highly variable, lasting from a few years to several decades before the onset of maturation (Tesch 2003). Upon maturation, the yellow eel transforms into silver eel and initiates its spawning migration back to the Sargasso Sea (Grassi 1897; Schmidt 1912, 1922; Miller et al., 2019; Wright et al., 2022). Given their life-history, the European eel was historically classified as catadromous, a classification that largely remains today (Schmidt 1912; 1922; Durif et al., 2023). However, it is well known that during the yellow eel life stage, individuals display great variation in habitat use; some eels remain in coastal areas with varying salinities and never enter freshwater, whereas others inhabit lakes and/or rivers, and some individuals shift between freshwater and coastal habitats one or several times before maturation (Tzeng et al., 1997; Limburg et al., 2003; Tesch, 2003; Daverat et al., 2006; Rohtla et al., 2023; Teichert et al., 2023). Despite the large body of scientific literature on the habitat use of eel, we have limited knowledge regarding the relative importance of coastal habitats in relation to freshwater systems concerning their contribution to the total production of silver eels. Such knowledge is crucial given the dramatic decline of the European eel population which is currently at a critically low level (ICES, 2025).

To investigate habitat choice of fish, otolith microchemistry is a well-established method that tracks changes in ambient water chemistry via the incorporation of trace elements into otolith material forming over the lifespan of an individual as it grows (Campana 1999; Campana & Thorrold 2001; Walther & Limburg, 2012). The ratio of strontium (Sr) to calcium (Ca) in the otoliths is positively correlated with salinity in the water (Kawakami et al. 1998; Marohn et al. 2009; Kraus & Secor 2004; Clément et al., 2014). Hence, variations in otolith Sr:Ca ratios in transects from the otolith core to the outer edge of the otolith provide a high-resolution trace-element diary revealing migration history and habitat use across salinity gradients throughout an individual’s lifetime.

Here, we used otolith microchemistry data from more than 3600 European eels from a large number of coastal and freshwater sampling sites across Sweden, and in coastal areas of Denmark, Finland, Estonia, and Poland, to assess individual habitat use. We analysed the Sr:Ca ratio of otolith transects, combined with established time-series clustering and segmentation methods, and thereby assigned life-histories to each individual eel in the categories: coastal resident, estuary resident, freshwater resident, and habitat shifters. This method allowed us to assess individual variation in habitat use of eel along a salinity gradient ranging from fully marine to freshwater.

## Material and Methods

All data handling, analysis and visualisation were performed using R v.4.3.1 (R Core Team, 2025) and the R-packages tidyverse v.2.0.0 (Wickham et al. 2019), DBI v.1.2.3 (R Special Interest Group on Databases et al., 2024), odbc v.1.4.2 (Hester et al., 2024), flextable v.0.9.5 (Gohel & Skintzos 2024), officer v.0.6.5 (Gohel & Moog 2024), webshot2 v.0.1.1 (Chang 2023), segclust2d v.0.3.3 (Patin et al. 2020), zoo v.1.8-12 (Zeileis & Grothendieck 2005), terra v.1.7-46 (Hijman 2023), sf v.1.0-15 (Pebesma 2018; Pebesma & Bivand 2023), viridis v.0.6.5 (Garnier et al., 2024), rnaturalearth v.1.0.1 (Massicotte & South 2023), ggrepel v.0.9.3 (Slowikowski 2023), ggnewscale v.0.5.2 (Campitelli 2025), ggspatial v.1.1.9 (Dunnington 2023), tidytext v.0.4.2 (Silge & Robinson 2016), patchwork v.1.3.0 (Pedersen 2024), ggtext v.0.1.2 (Wilke & Wiernik 2022) and ggmagnify v.0.4.1.9 (Hugh-Jones 2024). Artificial intelligence (OpenAI 2026) was used to assist with R-code troubleshooting, debugging, and optimization. All generated code were validated by the authors.

### Data origin

The otolith microchemistry data originated from eels sampled as part of various research projects and within several monitoring programmes (including the ongoing Swedish eel monitoring program within the EU Data Collection Framework, DCF). The data was accessed via the Institute of Freshwater Research, Sweden (formerly part of the Swedish Board of Fisheries, currently part of the Swedish University of Agricultural Science, SLU). The data had been collected over a period of 40 years, from 1984 to 2024. Since the purpose of collection varied, different types of catch methods had been used (e.g. fyke-nets, electrofishing, traps, eel collectors), and in some cases, eel had been sampled from the landed catch of commercial eel fishers. Following collection, eels were kept frozen (−20°C) and later thawed and dissected to obtain individual-level data, and otoliths were extracted and archived for subsequent use. Extracted otoliths have, for example, been used for age determination, shape-analysis (Fabosa, 2002), and for otolith microchemistry analyses (using different methods, see Otolith microchemistry data section below) to monitor the distribution of restocked eels in Swedish waters (since 2009, all restocked eels are chemically Sr-marked before being released; Wickström & Sjöberg, 2014). Most of the otolith microchemistry data have never been published, and the complete dataset has not been analysed as one unit.

### Study area

The data used in this study originated from eels sampled along the Swedish west– and east coasts, in coastal areas of Denmark, Finland, Estonia, and Poland, and in inland waters of Sweden (Fig. 1). As such, the main study area is the Skagerrak and Kattegat area, and the Baltic Sea region around Sweden. The Baltic Sea is one of the world’s largest brackish water bodies, connected to the North Sea via the Danish Straits (Little Belt, Great Belt, and Öresund), Kattegat, and Skagerrak. The salinity gradient in this region ranges from fully marine (>30 PSU) in the Skagerrak to nearly freshwater (<1 PSU) in the north of the Baltic Sea (Bothnian Bay) (Fig. 1) (Bonsdorff, 2006). In addition, there are more than 100 major watersheds with a drainage area larger than 200 km² in Sweden, spanning from the northernmost west coast (Skagerrak) to the northernmost east coast (Bothnian Bay, Baltic Sea) (SMHI, 2025). For naturally recruiting eel from the Sargasso Sea, which enter this region via Skagerrak and Kattegat, individuals have access to a large variety of potential growth habitats varying in salinity, making it a suitable study system for assessing variation in habitat use of eel.

**Figure 1.**
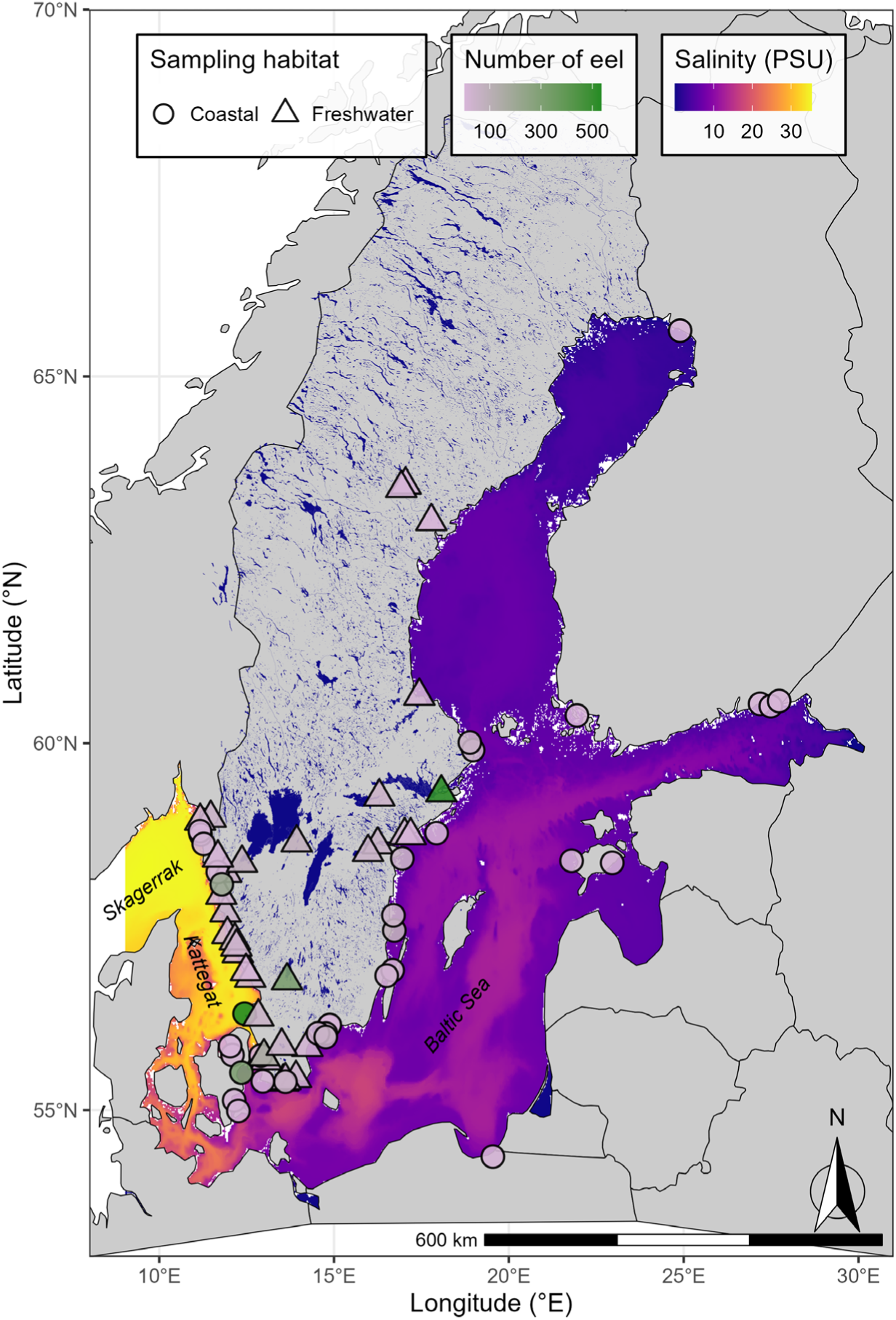
Map showing the study area consisting of Skagerrak and Kattegat (Sweden and Denmark), the Baltic Sea (Sweden, Denmark, Finland, Estonia, Poland), and freshwater sites (Sweden) with a colour gradient corresponding to sea floor salinity, with high salinity in yellow to low salinity in blue (practical salinity unit, PSU). Sampling sites are indicated with a circle for coastal sites and a triangle for freshwater sites. The sampling site symbols are filled using a colour gradient indicating the number of eels sampled per site (n = less than 100 in pink to n = more than 300 in green). Data on sea floor salinity was downloaded from the Copernicus Monitoring Environment Marine Service web portal (CMEMS 2026).

### Individual eel data

Individual-level eel data (i.e., the dissection data, see Data origin section above) were accessed from two databases hosted by SLU, Department of Aquatic Resources (SLU Aqua); Sötebasen (locally hosted by the Institute of Freshwater Research, SLU Aqua) and KUL, the database for fish monitoring along the coast (KUL 2026). In total, dissection data for 3648 individuals were extracted, of which 3439 were extracted from Sötebasen (extracted date: 2026-02-12), and 209 individuals were extracted from KUL (2026-02-12). Data on sampling site, sampling year, total body length (mm), total body weight (mass in gram) and individual ID were extracted from the databases and merged with the individual otolith microchemistry data (Table 1).

**Table 1.**
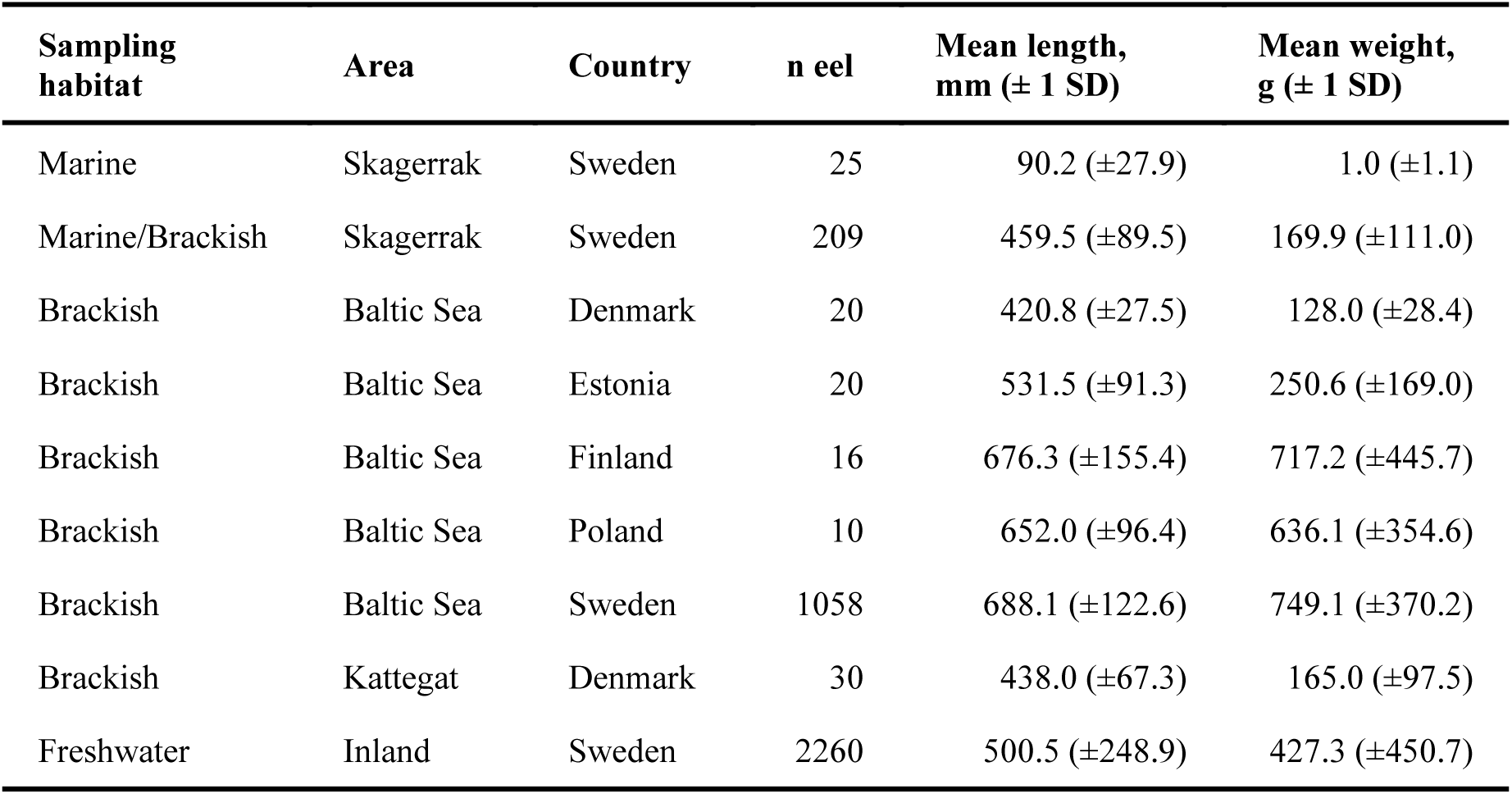
Summary of data extracted from the databases Sötebasen and KUL for dissected eel with existing otolith microchemistry data, detailing which habitat each eel had been captured and sampled from (marine, brackish, freshwater), the geographic area, the country, the number of eels, mean total body length (mm ± 1 standard deviation [SD]), and mean total body weight (g ± 1 SD).

| Sampling habitat | Area | Country | n eel | Mean length, mm (± 1 SD) | Mean weight, g (± 1 SD) |
| --- | --- | --- | --- | --- | --- |
| Marine | Skagerrak | Sweden | 25 | 90.2 (±27.9) | 1.0 (±1.1) |
| Marine/Brackish | Skagerrak | Sweden | 209 | 459.5 (±89.5) | 169.9 (±111.0) |
| Brackish | Baltic Sea | Denmark | 20 | 420.8 (±27.5) | 128.0 (±28.4) |
| Brackish | Baltic Sea | Estonia | 20 | 531.5 (±91.3) | 250.6 (±169.0) |
| Brackish | Baltic Sea | Finland | 16 | 676.3 (±155.4) | 717.2 (±445.7) |
| Brackish | Baltic Sea | Poland | 10 | 652.0 (±96.4) | 636.1 (±354.6) |
| Brackish | Baltic Sea | Sweden | 1058 | 688.1 (±122.6) | 749.1 (±370.2) |
| Brackish | Kattegat | Denmark | 30 | 438.0 (±67.3) | 165.0 (±97.5) |
| Freshwater | Inland | Sweden | 2260 | 500.5 (±248.9) | 427.3 (±450.7) |

### Otolith microchemistry data

Two methods had been used to obtain microchemical data from eel otoliths: 1) field emission electron probe microanalysis (FE-EPMA, n=1958) and 2) laser ablation inductively coupled plasma mass spectrometry (LA-ICP-MS, n= 1690). Prior to being analysed using either method, the sagittal otolith had been embedded in epoxy (EpoFix, Struers) together with a small paper note with the ID-number of the individual eel, left over-night to be fixated, and then grinded from the sulcus side with a grinding machine (Buehler Phoenix Beta Grinder/Polisher, Buehler MetaServ® 250 Grinder-Polisher) using successively finer paper (P2500-4000) until the otolith core was exposed. The otolith had then been polished using a polishing cloth (MicroCloth) attached to the grinding machine and a polishing agent (Buehler MasterPrep, Polishing Suspension 0.05 µm).

#### Field emission electron probe microanalysis, FE-EPMA

The otolith microchemistry analyses using FE-EPMA had been conducted at the Department of Earth Sciences, Uppsala University, Sweden. The FE-EPMA was equipped with five wavelength dispersive spectrometers (WDS) as well as backscattered electron detectors (BSE). The field emission microprobe scans the sample surface and creates two-dimensional mappings that reveal differences in the chemical composition of the otoliths (Fig. 2A). After having placed the epoxy-embedded, polished otoliths on glass slides (16 otoliths/slide), the otoliths had been photographed, and a transect had been marked on the digital images. The transect had been laid out as far as possible from the core to the edge posteriorly (Limburg et al., 2003; Clevestam & Wickström, 2008), sometimes with adjustments to avoid cracks and possible vaterite inclusions crossing the measurement transect. Prior to analysis, all otoliths had been carbon-coated. The microprobe used a voltage of 20 kV and a current of 20 nA. The diameter of the electron beam was 15 µm. Strontianite (SrCO3) and calcite (CaCO3) were used as calibration standards. Strontium and calcium were measured at 30 measurement points along the previously determined transect. The distances between the measurement points were kept constant for each otolith but varied depending on the otolith size (Fig. 2A). The Sr:Ca ratio was derived by dividing Sr/Ca * 1000 (wt%).

**Figure 2.**
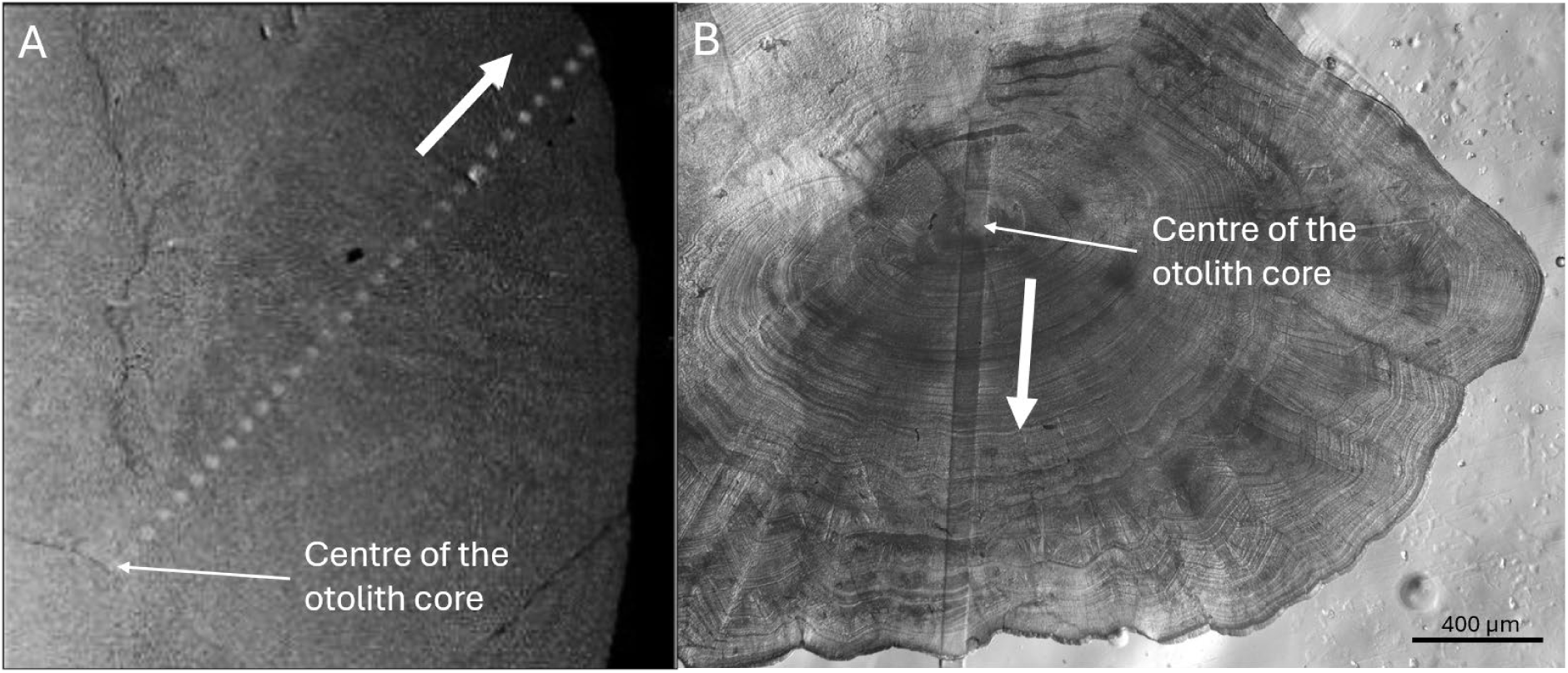
Example photos showing the otolith microchemistry transect analyses using. **A)** field emission electron probe microanalysis, FE-EPMA or **B)** laser ablation inductively coupled plasma mass spectrometry, LA-ICP-MS. White arrows show the direction of the transect, extending from the otolith core to the otolith edge. In panel **A,** the transect begins at the centre of the otolith core and consists of 30 datapoints (grey dots). In panel **B,** the transect begins in the outer region of the otolith core and consists of overlapping data points that form a continuous transect through the centre of the core, visible as a rectangular trench on the otolith surface following analysis.

#### Laser ablation inductively coupled plasma mass spectrometry, LA-ICP-MS

The otolith chemistry analyses using LA-ICP-MS had been conducted at the Department of Geology, Lund University, Sweden. Prior to analysis, the epoxy-embedded, polished otoliths had been photographed and a proposed transect had been marked on the digital image; the otoliths had then been mounted on glass slides (16 otoliths/slide). The laser followed the proposed transect through the core and out to the edge of the otolith (Fig. 2B). To avoid contamination, pre-ablation was performed prior to analysis. Helium served as the carrier gas, with additional argon introduced downstream of the sample chamber. The laser spot size ranged from 30 x 70 µm to 40 x 90 µm, due to several adjustments over the years and has thus varied over time. The scanning speed has also varied over time, ranging from 6 µm s^-1^ to 32 µm s^-1^. The most frequently used configuration was 30 x 70 µm with a scan speed of 9 µm s^-1^. For the quantification of Sr and Ca, the reference material MACS-3 was used as the primary standard, glass NIST SRM 610 and 612 as the secondary standards and ^43^Ca as the internal standard assuming an otolith Ca content of 38 wt% in a standard-sample bracketing setup. Data reduction, background subtraction, correction of instrument drift and the calibration of trace elemental concentrations in parts per million (ppm) was done, using the Iolite v3 within Igor Pro software (Hellström et al., 2008; Paton et al., 2011). As ^43^Ca was used as the internal standard, ^43^Ca counts per second (CPS) values were converted to ppm by multiplying the CPS with 380000 and then divide it by the mean CPS for each individual. To remove outliers and Ca-values corresponding to the epoxy and not the otolith, all Ca-values ±1.25 SD of the mean Ca-value of each otolith transect profile were removed. Sr:Ca ratios are expressed as (Sr concentration [ppm] *1000 / Ca concentration [ppm]). Detailed information about the LA-ICP-MS instrument settings is summarised in the supplementary information (SI, Table S1).

### Determination of Sr:Ca habitat-specific threshold values using reference eel

To assign habitat use based on salinity, Sr:Ca ratios were used as a proxy, and threshold values were established using reference eels. The reference eels were naturally recruited eels from two freshwater sites for FE-EPMA otolith chemistry data (n=27) and from a gradient of freshwater and coastal sites for LA-ICP-MS otolith chemistry data (n=77) (SI, Table S2). Threshold values were determined for FE-EPMA and LA-ICP-MS separately, since the methods to calculate the concentration of elements differ (wt% for FE-EPMA and ppm for LA-ICP-MS) and the amount of data-points per transect differed, with FE-EPMA having a fixed amount of data-points (n=30), while LA-ICP-MS generates more datapoints which vary between individuals depending on the otolith size (SI, Fig. S2, S3). Habitat-specific threshold values in Sr:Ca ratios were derived following the methodology described by Denis et al. (2023) and Teichert et al. (2023), using the R-package segclust2d developed for performing segmentation and clustering of time-series data in R (Patin et al., 2020) (Fig. 3). First, the otolith transect was divided into segments using the *segclust()* function (Fig. 3A, 3B). For FE-EPMA data, segmentation was performed with a minimum segment length of five data points and a maximum of six segments (Kmax=6). For LA-ICP-MS data, segmentation was performed with a minimum segment length of ten data points and a maximum of twenty segments (Kmax = 20). The number of clusters was evaluated for values between two and five, with the optimal model selected by the algorithm, determined based on the Bayesian information criterion (BIC). Secondly, the mean Sr:Ca ratio and standard deviation were calculated for each cluster within each individual transect. The resulting cluster means from all the reference eels were then used to determine threshold values for the clusters (Fig 3C, 3D). Each cluster was then interpreted and translated into a habitat salinity based on the known birthplace of European eel (the Sargasso Sea, fully marine), the known capture site for each individual (corresponding to the outermost region of the otolith), and the needed migration from the birthplace to reach each capture site (segments between the otolith core and the outer most segments of the otolith transect) (Tzeng et al., 1997) (Fig. 3E, 3F). For eel with otolith chemistry data from FE-EPMA analyses, the most common clustering of the reference eel transects, according to BIC, was a three-cluster classification of the Sr:Ca ratios (Fig. 3A). These clusters correspond to freshwater, coastal/brackish, and marine habitats with no overlap in mean Sr:Ca values (Table 2, Fig. 3A, 3C, 3E). For eel with otolith chemistry data from LA-ICP-MS analyses, the most common clustering of the reference eel transects, according to BIC, was a five-cluster classification based on Sr:Ca ratios (Fig. 3B). These clusters correspond to freshwater, estuarine, coastal/brackish and marine habitats (Table 2, Fig. 3B, 3D, 3F). Due to some overlapping outliers of the mean Sr:Ca between clusters derived from LA-ICP-MS data, mean Sr:Ca values within the 5^th^ and 95^th^ percentile range was used to determine Sr:Ca threshold values for each habitat type (Table 2, Fig. 3D, 3F).

**Figure 3.**
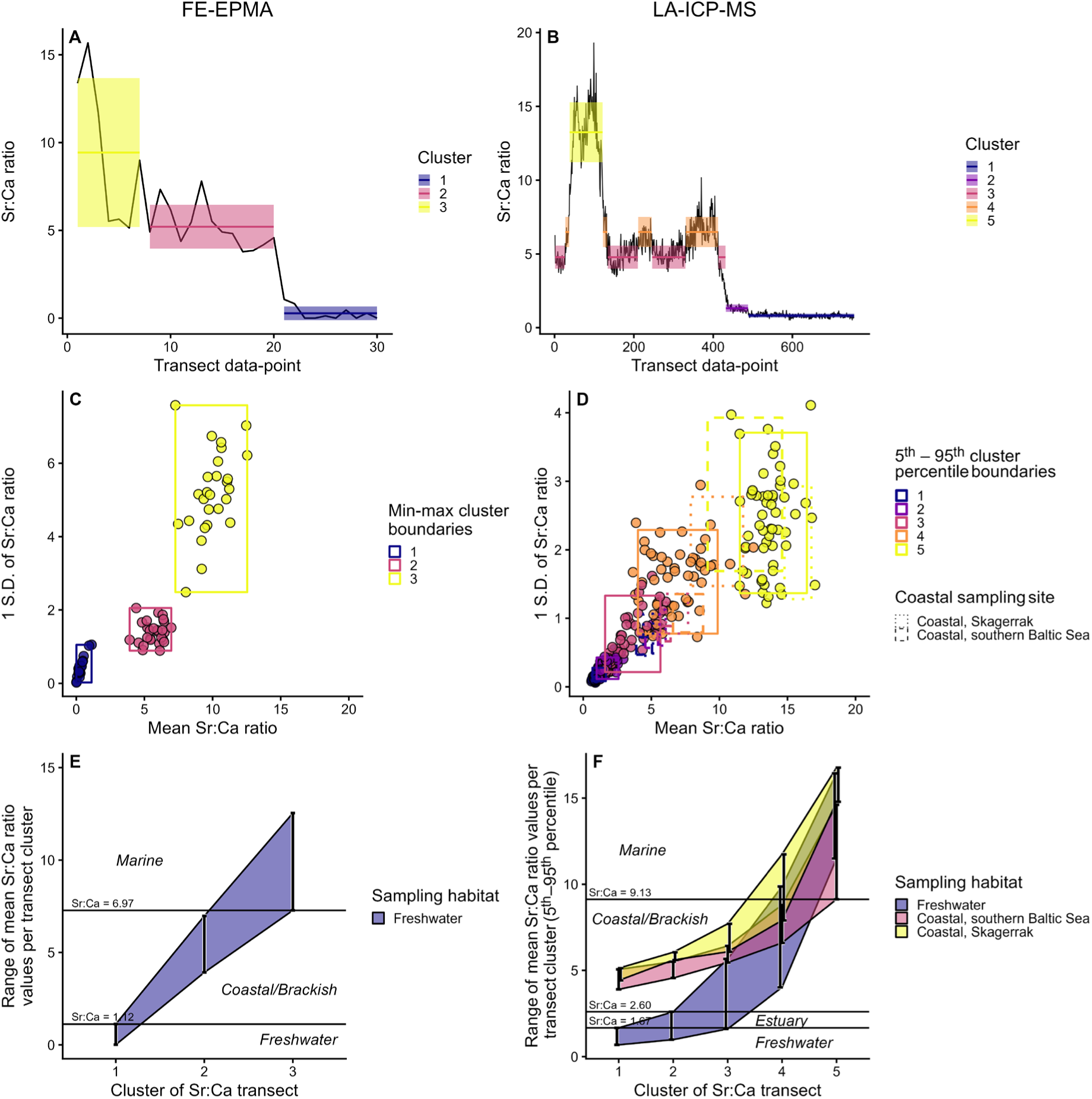
Strontium to calcium ratio (Sr:Ca) values obtained from eel otolith microchemistry transects, as well as the derived segments and clusters for data acquired using field emission electron probe microanalysis, FE-EPMA (panels **A, C, E**), and laser ablation inductively coupled plasma mass spectrometry, LA-ICP-MS (panels **B, D, F**). Panels **A** and **B** show the Sr:Ca ratio transects, including derived segmentation and clustering with cluster mean and SD (coloured horizontal solid lines with associated ribbon) for two individual eels caught in Lake Mälaren (freshwater lake) as examples of A) FE-EPMA and B) LA-ICP-MS results. Panels **C and D** show the relationship between cluster mean Sr:Ca ratios (x-axis) and their associated variation (± 1 SD, y-axis) for all reference eels. Each datapoint represents the Sr:Ca value for a given cluster within an individual eel. Boxes show the range of cluster mean Sr:Ca ratios, represented as the full range in panel **C** and the 5^th^– 95^th^ percentile range in panel **D**. In panel **D,** solid-line boxes represent eels sampled at freshwater sites, whereas dotted and dashed boxes represent eels caught at coastal sites (dotted = Kattegat, Stenungsund; dashed = southern Baltic Sea, Karlshamn). Panels **E** and **F** show the range of mean Sr:Ca ratios for each derived cluster, indicated by vertical lines and shaded ribbons, divided by capture habitat; Freshwater (purple), Coastal, southern Baltic Sea (pink) or Coastal, Skagerrak (yellow). Horizonal black lines indicate the derived Sr:Ca threshold values corresponding to growth in habitats with different salinities.

**Table 2.** Derived Sr:Ca threshold values from reference eels, corresponding to different growth habitat salinities based on either field emission electron probe microanalysis, FE-EPMA or laser ablation inductively coupled plasma mass spectrometry, LA-ICP-MS otolith microchemistry analyses.

| Growth habitat | Sr:Ca threshold values | Analytical method |
| --- | --- | --- |
| Freshwater | 0 - 1.12 | FE-EPMA |
| Coastal/Brackish | >1.12 - 6.97 | FE-EPMA |
| Marine | >6.97 | FE-EPMA |
| Freshwater | 0 - 1.67 | LA-ICP-MS |
| Estuary | >1.67 - 2.60 | LA-ICP-MS |
| Coastal/Brackish | >2.60 - 9.13 | LA-ICP-MS |
| Marine | >9.13 | LA-ICP-MS |

### Conversion of Sr:Ca ratios to habitat use and life-history assignment

The derived Sr:Ca ratio threshold values from the reference eels, corresponding to different habitats, were used to convert Sr:Ca transect ratios to habitat use over ontogeny for each individual eel in the dataset. After each Sr:Ca datapoint was converted to a corresponding habitat (Table 2), each individual was assigned a life-history, being either freshwater resident, estuary resident, freshwater/estuary resident, coastal resident or a habitat shifter (Table 3, Fig. 4). Since all eels are initially arriving to coastal areas from the Sargasso Sea, the initial coastal Sr:Ca values in the transect, corresponding to the arrival from the Sargasso Sea (e.g. Arai et al. 2006), were excluded from the life-history assignment to avoid classifying all eels as habitat shifters. Life-history assignment was only conducted for eel with a clear otolith core since this was needed to identify the centre of the core. Otoliths (and hence eel individuals) without a clear core were excluded. Individuals under 20 cm in length (n=523) were also excluded in subsequent analyses to ensure that all included eels had sufficient time to make an active habitat choice following their arrival from the Sargasso Sea.

**Figure 4.**
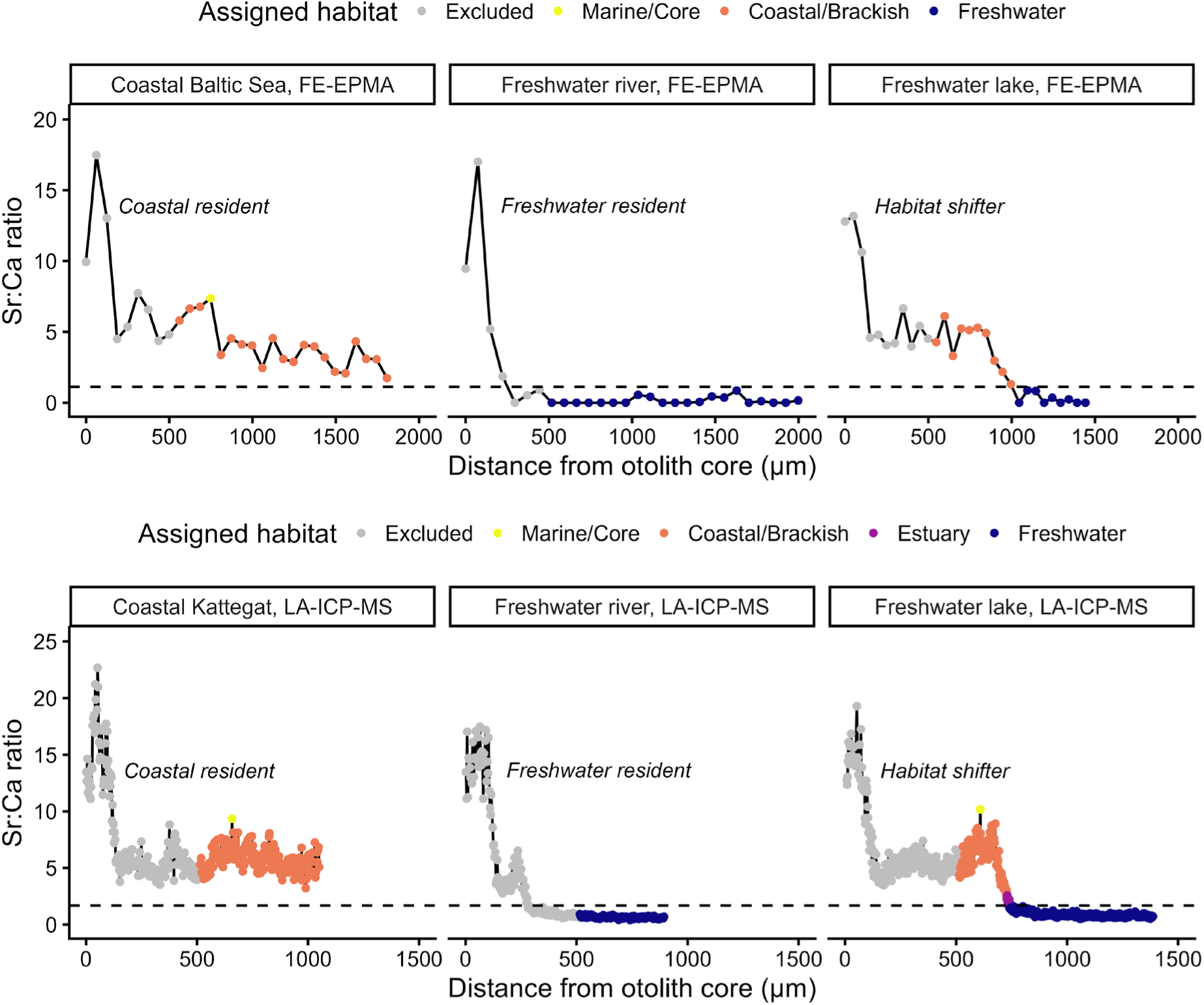
Examples of individual otolith microchemistry transects derived using either FE-EPMA (top panels) or LA-ICP-MS (lower panels). Each Sr:Ca data point has been converted to a corresponding habitat according to the derived Sr:Ca ratio threshold values (colour of each data-point) and each individual have been assigned with a life-history (written in italics in each panel). Panel titles show the sampling habitat of each individual and the analytical method used. Horizontal dashed lines show the derived threshold value corresponding to freshwater for FE-EPMA (Sr:Ca=1.12) and LA-ICP-MS (Sr:Ca=1.67).

**Table 3.**
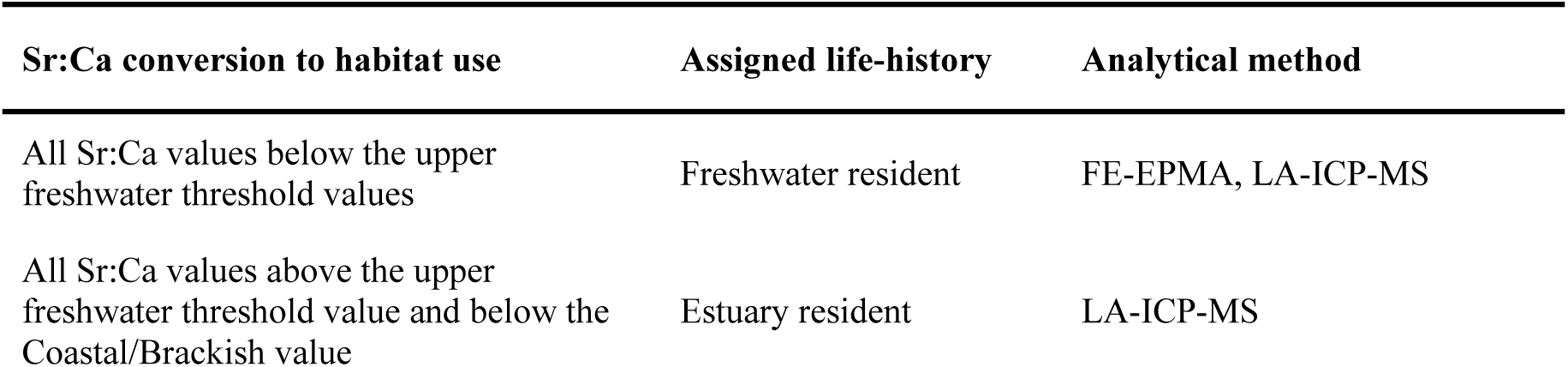
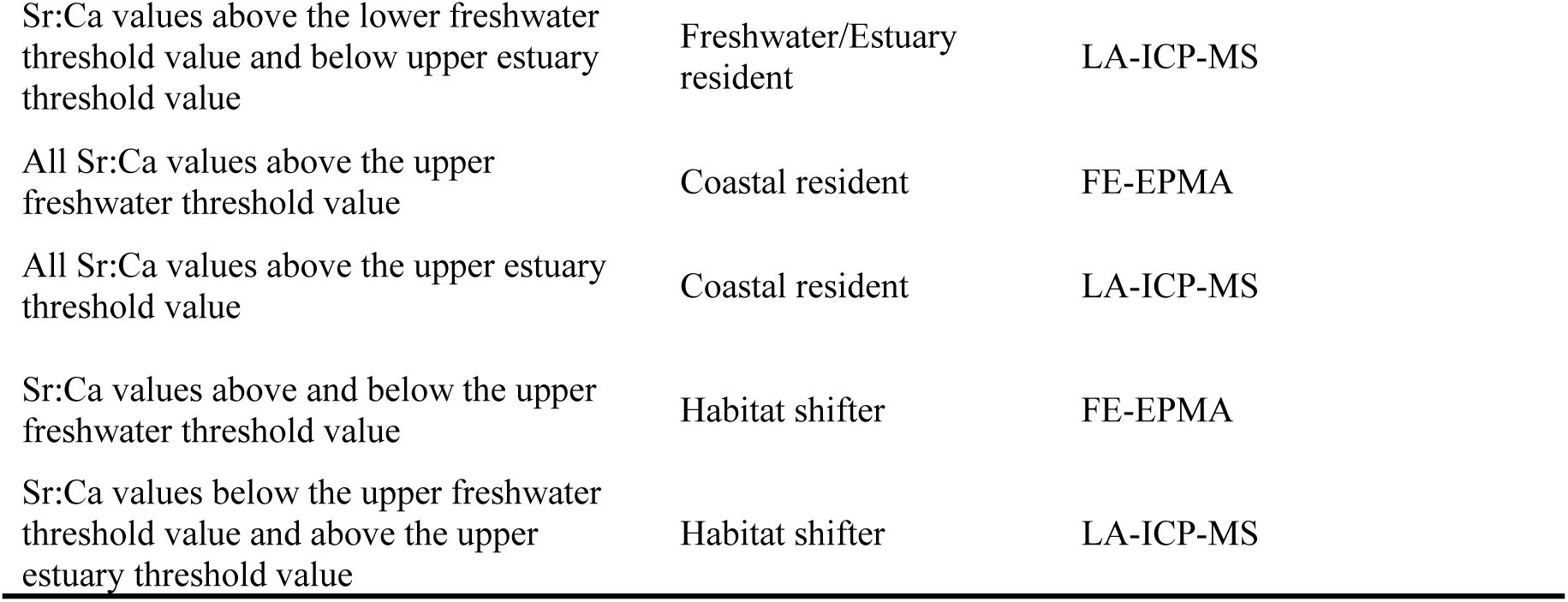
Conversion of Sr:Ca ratios to habitat use and subsequent life-history assignment.

For eel with LA-ICP-MS otolith chemistry data, the Sr:Ca conversion to habitat use and the subsequent life-history assignment was done accordingly:

1. Sr:Ca data points corresponding to the Sargasso Sea (i.e. the otolith core) were identified using the derived threshold value of the reference eels (Sr:Ca >9.13, Table 2).
2. The number of data points for each contiguous Sr:Ca region identified as the otolith core was quantified for each individual, applying an 8 data-point window smoother using the *rollmean()* function of the R-package zoo to ignore the influence of single-points with values <9.13 in Sr:Ca.
3. Individuals without a distinct otolith core signal in their Sr:Ca transect, corresponding to a contiguous region with Sr:Ca >9.13 longer than 20 consecutive data-points, were removed (lack of core probably due to over-grinding in the otolith preparation process).
4. The longest contiguous region corresponding to the otolith core was selected, and in the case of equally long regions, the region with highest Sr:Ca value was selected to represent the otolith core.
5. The mid-point of the selected region from step 3 and step 4 was set to zero, representing the centre of the otolith core. Then, the distance from the zero-point was calculated for each data-point and transect by multiplying the elapsed scanning time (seconds) with the scanning speed (µm s^-1^) of the laser.
6. The mean distance from the core to the otolith edge was calculated for all eels between 10-20 cm in length and was used to exclude the initial coastal Sr:Ca signal corresponding to the initial arrival of eel to the coast from the Sargasso Sea (n= 348, mean distance from zero-point of the otolith core to the otolith edge = 513 µm).
7. Conversion of Sr:Ca ratio to habitat use (Table 2) was conducted for all data points in each transect starting from 513 micrometres distance from the otolith core, while individuals with a shorter transect than 513 micrometres from the otolith core were removed (n=121).

For eel with FE-EPMA otolith chemistry data, the distance between each consecutive data-point along the transect was calculated using the Euclidian distance using the X and Y coordinates for each measurement point according to equation 1:

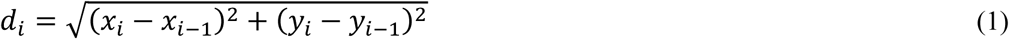

where *dᵢ* represents the distance between consecutive points *i* and *i*−1, and *x* and *y* denote their respective Cartesian coordinates. The cumulative distance was then calculated for all data points along the transect to get the distance from the centre of the core for each data point. After the distance from the otolith core was calculated for each data-point, conversion of the Sr:Ca ratio to habitat use (Table 2) was conducted for all data points in each transect starting from 513 micrometres distance from the otolith core, while individuals <20 cm in length (n=92) or with a shorter transect than 513 micrometres from the otolith core were removed (n=0).

Eels fulfilling the above criteria (n_total_=2938; n_LA-ICP-MS_=1045, n_FE-EPMA_=1893), were classified as either freshwater resident, estuary resident, coastal resident, or habitat shifter (Table 3, Fig. 4). For eels classified as habitat shifters, the number of shifts between freshwater and coastal/brackish habitats was quantified.

## Results

For eels caught at coastal sites, the majority were classified as coastal residents (79-99% for eels captured at Skagerrak/Kattegat sites; 85-100% for eels captured at Baltic Sea sites, for both otolith chemistry methods) (Table 4). For eels caught at freshwater sites draining to Skagerrak and Kattegat, the majority were classified into two main life histories; freshwater residents (58%), followed by habitat shifters (43%) for eel with FE-EPMA otolith chemistry data, while for eel with LA-ICP-MS otolith chemistry data there were three main life histories; freshwater residents (51%) followed by freshwater/estuary residents (27%) and habitat shifters (19%) (Table 4). For eels caught at freshwater sites draining to the Baltic Sea, the majority was classified as freshwater residents (63%), followed by habitat shifters (36%) for eel with FE-EPMA otolith chemistry data, while freshwater residents was the most common life history (45%), followed by habitat shifters (26%) and freshwater/estuary residents (14%) for eels with LA-ICP-MS otolith chemistry data (Table 4). There was large variation in the proportion of the different life-histories among sampling sites within each major geographical sampling region (Fig. 5).

**Figure 5.**
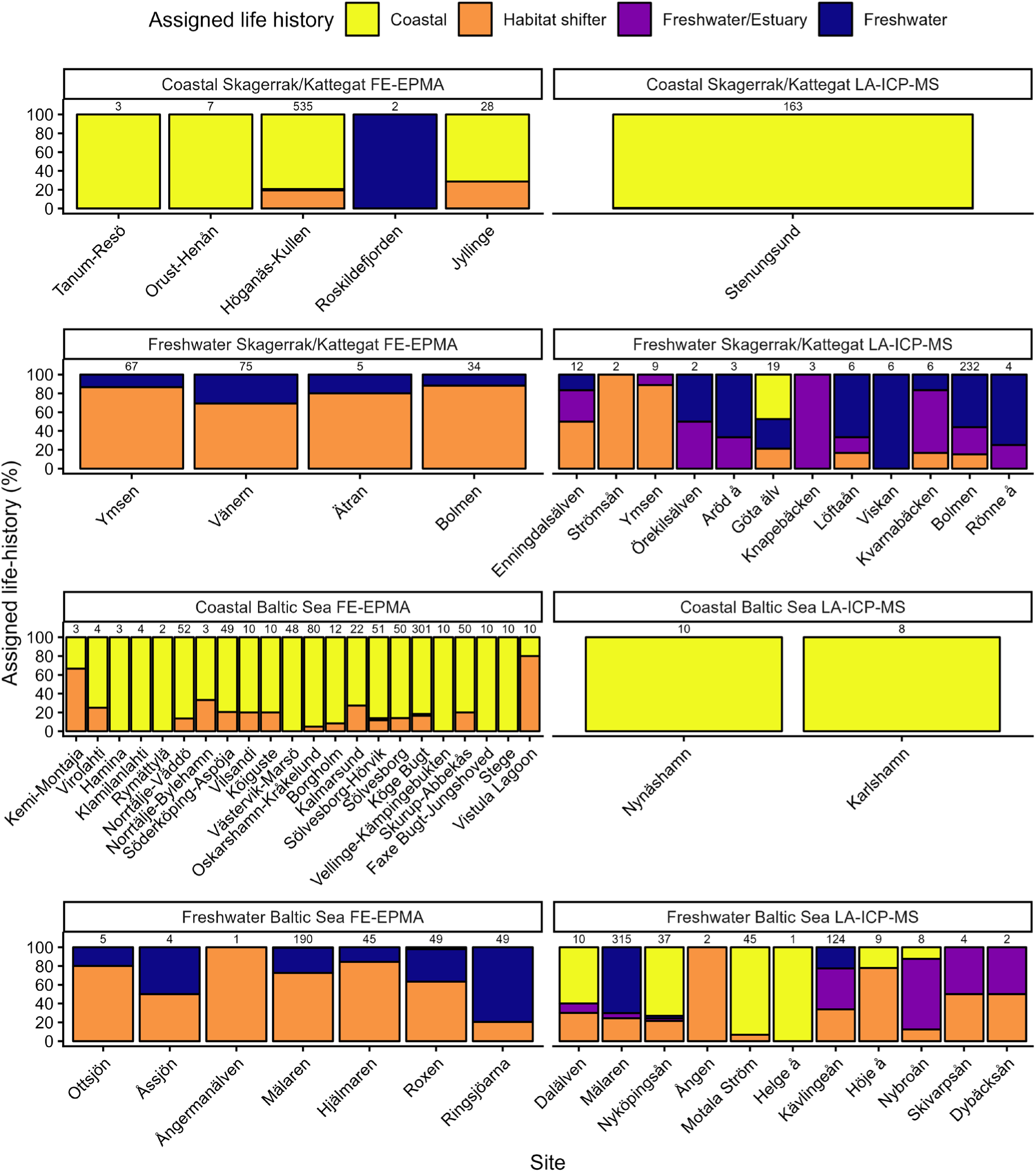
Percentage of eels per assigned life-history category for each sampling site per geographical area, sorted from north to south within each panel, divided by otolith microchemistry method (field emission electron probe microanalysis, FE-EPMA, or laser ablation inductively coupled plasma mass spectrometry, LA-ICP-MS). The total number of individuals per sampling site is shown above each bar.

**Table 4.** Number and percentage (out of the total sample size) of individuals per assigned life history (coastal resident, freshwater resident, estuary resident, freshwater and estuarine and habitat shifter) based on Sr:Ca transect values for eels sampled at coastal sites in Skagerrak/Kattegat and the Baltic Sea, and eels sampled at freshwater sites draining to either Skagerrak/Kattegat or the Baltic Sea, based on otolith microchemistry analyses (analytical method: field emission electron probe microanalysis, FE-EPMA, or laser ablation inductively coupled plasma mass spectrometry, LA-ICP-MS). Assigned life histories marked with an asterisk indicate incorrectly assigned (sampled at freshwater sites but assigned as coastal residents).

| Sampling habitat | Sampling region | Assigned life history | Total N | Assigned n | Assigned % | Analytical method |
| --- | --- | --- | --- | --- | --- | --- |
| Coastal | Skagerrak/Kattegat | Coastal resident | 575 | 455 | 79.1 | FE-EPMA |
| Coastal | Skagerrak/Kattegat | Freshwater resident | 575 | 9 | 1.6 | FE-EPMA |
| Coastal | Skagerrak/Kattegat | Habitat shifter | 575 | 111 | 19.3 | FE-EPMA |
| Coastal | Skagerrak/Kattegat | Coastal resident | 163 | 162 | 99.4 | LA-ICP-MS |
| Coastal | Skagerrak/Kattegat | Habitat shifter | 163 | 1 | 0.6 | LA-ICP-MS |
| Freshwater | Skagerrak/Kattegat | Freshwater resident | 181 | 37 | 20.4 | FE-EPMA |
| Freshwater | Skagerrak/Kattegat | Habitat shifter | 181 | 144 | 79.6 | FE-EPMA |
| Freshwater | Skagerrak/Kattegat | Coastal resident* | 304 | 10 | 3.3 | LA-ICP-MS |
| Freshwater | Skagerrak/Kattegat | Freshwater resident | 304 | 154 | 50.7 | LA-ICP-MS |
| Freshwater | Skagerrak/Kattegat | Freshwater/Estuary | 304 | 83 | 27.3 | LA-ICP-MS |
| Freshwater | Skagerrak/Kattegat | Habitat shifter | 304 | 57 | 18.8 | LA-ICP-MS |
| Coastal | Baltic Sea | Coastal resident | 794 | 671 | 84.5 | FE-EPMA |
| Coastal | Baltic Sea | Freshwater resident | 794 | 6 | 0.8 | FE-EPMA |
| Coastal | Baltic Sea | Habitat shifter | 794 | 117 | 14.7 | FE-EPMA |
| Coastal | Baltic Sea | Coastal resident | 21 | 21 | 100.0 | LA-ICP-MS |
| Freshwater | Baltic Sea | Coastal resident* | 343 | 2 | 0.6 | FE-EPMA |
| Freshwater | Baltic Sea | Freshwater resident | 343 | 117 | 34.1 | FE-EPMA |
| Freshwater | Baltic Sea | Habitat shifter | 343 | 224 | 65.3 | FE-EPMA |
| Freshwater | Baltic Sea | Coastal resident* | 557 | 79 | 14.2 | LA-ICP-MS |
| Freshwater | Baltic Sea | Freshwater resident | 557 | 250 | 44.9 | LA-ICP-MS |
| Freshwater | Baltic Sea | Freshwater/Estuary | 557 | 82 | 14.7 | LA-ICP-MS |
| Freshwater | Baltic Sea | Habitat shifter | 557 | 146 | 26.2 | LA-ICP-MS |

A small proportion of eels were assigned with the wrong life-history, with 3% of the eels caught in freshwater sites draining to Skagerrak/Kattegat being incorrectly classified as coastal residents, and 0.6% (FE-EPMA) and 14% (LA-ICP-MS) of the eels caught in freshwater sites draining to the Baltic Sea being incorrectly classified as coastal residents (Table 4). The latter category (eels caught in freshwater sites draining to the Baltic Sea) mainly consisted of individuals from two sampling sites; river Nyköpingsån and river Motala Ström (Fig. 5).

### Habitat shifts

For the 596 eel with FE-EPMA chemistry data assigned with a habitat-shifting life-history, the number of shifts between coastal and freshwater habitats varied between 1-14 shifts, with the majority of the individuals shifting once (n=245, 41%), followed by two and three shifts (n=132, 22%; n=91, 15% respectively) (Fig. 6A). For the 204 eel with LA-ICP-MS chemistry data assigned with a habitat-shifting life-history, the number of shifts between freshwater/estuary and coastal habitats varied between 1-59 shifts, with the majority shifting twice (n=96, 47%), followed by one and three shifts (n=33, 16%; n=17, 8% respectively) (Fig. 6B).

**Figure 6.**
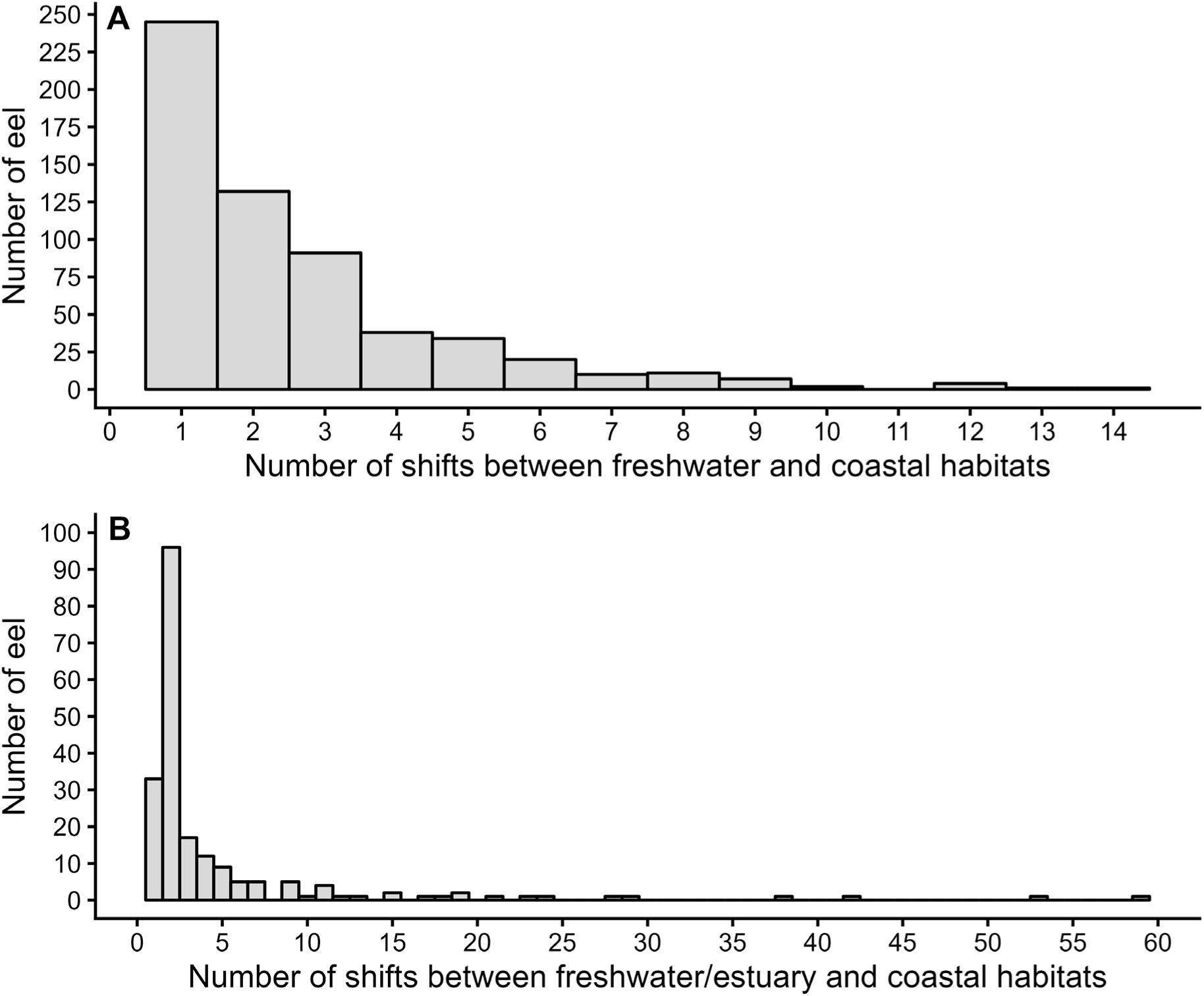
Number of eels assigned with a habitat-shifting life-history that. **A** had shifted 1-14 times between freshwater and coastal habitats based on FE-EPMA otolith microchemistry data, **B** had shifted 1-59 times between freshwater/estuary and coastal habitats based on LA-ICP-MS otolith microchemistry data.

## Discussion

Here, using existing otolith microchemistry data of more than 3600 eels caught along a salinity gradient from fully marine to freshwater, we show that eels display large variation in habitat use during their growth phase in the Skagerrak, Kattegat and Baltic Sea region. We found eels that had been resident on the coast, in freshwater, as well as eels using both freshwater/estuaries and coastal habitats after their arrival from the Sargasso Sea. For individuals using both freshwater/estuarine and coastal habitats (habitat shifters), the majority shifted between habitats 1-3 times. Our findings corroborate previous studies categorizing eels as a facultative catadromous fish species, showing high intraspecific variability in habitat use, adding complexity to the already diverse and complex life-history of the European eel.

That eel uses coastal habitats, and not just freshwater habitats, to grow has been observed in several regions of its distribution area (Daverat et al 2006; Durif et al. 2023), including the Mediterranean Sea (Capoccioni et al., 2013; Tahri & Panfili, 2023; Teichert et al., 2023), British Isles (Arai et al., 2006, Teichert et al., 2023), Norwegian coast (Rothla et al., 2023) and at the north Atlantic coast of France (Daverat & Tomás, 2006). We found that the majority of the eels caught at coastal sites used only coastal habitats for growth, both in Skagerrak, Kattegat and the Baltic Sea. These findings strengthen other indications of long-term existence of eels growing at the coast in the region, e.g. the long history of commercial fisheries targeting yellow eel on the Swedish west coast (Magnusson & Dekker, 2021) and the increase and stabilization of the yellow eel standing stock and estimated silver eel production on the Swedish west coast following a complete eel fishing ban in 2012 (Säterberg et al., 2025). Note that this does not necessarily imply that the eels had never visited a freshwater site, but it means that they had not spent enough time in freshwater to grow, and hence no freshwater signal had been incorporated in their otoliths (further discussed below). Relevant to this aspect is also the experimental observation that eels may grow slower in freshwater compared to saltwater, despite identical food supply (Édeline & Élie 2004). We further show that a large proportion of eels caught at the straits between the Baltic Sea and Kattegat (i.e. silver eels migrating towards the Sargasso Sea) had been coastal residents (Fig. 5). Given that current assessments of the status of the European eel are largely based on recruitment to freshwater systems (Pike et al., 2020; ICES, 2025), these findings highlight the potential need for developing a more regional framework for future eel assessments which also incorporates recruitment, growth, and production of silver eels in coastal systems.

Freshwater ecosystems have been heavily impacted by human activities. Hydroelectric power production, ditching, urbanization, land use change, and flood-control infrastructure have altered natural flow regimes, reduced connectivity through fragmentation and migration barriers, and decreased the survival of species passing through hydropower turbines, negatively affecting many fish species (Nilsson et al., 2005; Poff et al. 2007; Darwall et al., 2018; Duarte et al., 2021), including eel (Limburg & Waldman, 2009; Calles et al. 2010; Tamario et al., 2019; Waldman & Quinn, 2022). Owing to the decreased accessibility and availability of freshwater habitats, one could question whether the coastal and marine residency of eel is more common in recent times compared to the pre-industrial era. According to Guiry & Robson (2024), eels used coastal sea grass meadows during the Late Mesolithic and Early-Middle Neolithic periods (∼5400 – 2550 calibrated years before common era (BCE)), based on stable isotope analyses of eel otoliths found at excavation sites in Denmark. In addition, Kettle et al. (2008) showed that historical eel remains have been found across much of the specieś current distribution area, including both coastal and inland areas. This suggests that the use of coastal habitats, as well as the high intraspecific variability in habitat use associated with eel growth, has also occurred historically. At the same time, eels have the capacity to climb vertical structures in air (Linton et al., 2007) and can survive for long time-periods on land (Berg & Steen, 1965). Such behaviours have likely evolved to enable eels to colonize remote or partially inaccessible freshwater habitats where they can feed and grow large (Ibbotson et al., 2002). Currently, eels that grow in freshwater habitats have a higher risk of mortality when leaving the system for migration towards the Sargasso Sea compared to eels growing in coastal areas, as the anthropogenic mortality of eels in freshwater systems has increased during the last 100 years due to hydropower development (e.g. Calles et al. 2010). This increased anthropogenic mortality has potentially decreased the number of eels from freshwater systems that survives the initial phase of their spawning migration. This could be a reason for the high proportion of coastal residential eels observed at coastal sampling sites. It is also possible that the eels residing along the coast tried to, but failed, to access freshwater sites due to the migration barriers, i.e., the coastal habitat is not chosen/not the preferred choice but the only available habitat for them. Whether the proportion of individuals not entering freshwater nor shifting between freshwater and coastal habitats have increased in recent time due to decreased freshwater habitat availability and/or decreased survival in freshwater systems due to anthropogenic activities is hard to disentangle, and more research is needed to assess individual variation in habitat use of eel during pre-industrial conditions.

Almost 30% of the eel in the total dataset were assigned as habitat shifters (n=800), with the majority of individuals shifting between habitats one to three times. The maximum number of shifts recorded was 14 with the FE-EMPA method and a remarkable 59 times with the LA-ICP-MS method. The greater number of shifts detected with LA-ICP-MS is likely due to the overlapping data points forming the continuous transect, as compared to the 30 data points from the FE-EMPA. Hence, the former method has a greater capacity to detect habitat shifts than the latter. It is therefore possible that rapid shifts taking place during little growth of the otolith is missed by the FE-EMPA method. This was confirmed by the method comparison, where 25 otoliths were analysed using both methods. While both methods assigned life history similarly (all assigned as habitat shifters, SI, Table S3), the number of habitat shifts assigned sometimes varied greatly, with the largest difference being one shift assigned using FE-EMPA while 15 habitat shifts could be detected with LA-ICP-MS for that very same individual (SI Table S3). Only seven of the 25 individuals in this comparison had the same number of detected habitat shifts (SI Table S3). This does not necessarily imply that FE-EMPS is an erroneous method to accurately assign habitat shifters, but if the aim is to study number of habitat shifts in detail, then the more precise LA-ICP-MS method should preferably be used.

Otolith microchemistry is a commonly used tool to infer and backtrack the salinity of habitats used by fish over ontogeny. Using the Sr:Ca ratio to distinguish between freshwater, brackish, and marine habitats and quantify shifts between habitats depends on determined threshold values for each habitat-type, varying between study areas and analytical methodologies (Table 2, Fig. 2, SI Fig. S1 & S2; Tzeng et al., 1997; Limburg et al., 2003; Marohn et al., 2011; Rothla et al., 2023; Denis et al., 2023; Teichert et al., 2023). Therefore, derived threshold values in Sr:Ca corresponding to a specific salinity are not universal and may require regional adjustment due to local water chemistry, sediment composition, and hydrological gradients (Hamer et al., 2015). Here, we used eels from selected reference freshwater and coastal sampling sites to derive method– and site-specific threshold values in Sr:Ca for growth in habitats with varying salinities. The freshwater classification was not 100% correct for either the FE-EPMA or the LA-ICP-MS analysed individuals, with the majority of the incorrectly classified eels caught at two adjacent freshwater sites (rivers): Motala Ström and Nyköpingsån, with LA-ICP-MS otolith microchemistry data. Why these individuals have been incorrectly classified could be due to several reasons. Most of the otolith preparations after 2009 have been done to enable the detection of the large-scale Sr-marking of restocked eel, evident as a circle with highly elevated Sr concentration around the otolith core (Wickström & Sjöberg, 2014). Thus, there is a possibility that exposing the core but avoiding “overgrinding” the otolith and the Sr-mark at the cost of not exposing the outermost regions of the embedded otoliths could have been prioritized. If so, the outermost region of the otolith, corresponding to the most recent growth habitat, would be lost. In addition, Sr accumulation in otoliths requires growth, and some eels might only have spent a short time in the freshwater system before being captured, while others have stayed longer, creating a mismatch between the final growth habitat and the sampling site. If this is the case, then the incorrectly assigned individuals is not an error in the method as such, but a consequence of the eel having spent too little time in freshwater for that signal to be incorporated (or only having spent time there in the winter when growth is heavily reduced). A third reason could be due to differences in Sr-levels in the water between watersheds, which affects the Sr:Ca ratio in the otolith also in freshwater (e.g. Rothla et al., 2023), which we cannot control for in our study as we do not have the water Sr-concentration from the sampling sites. Site-specific concentrations of various elements from water samples collected at the sampling sites would be ideal to increase the precision of the derived threshold values in Sr:Ca corresponding to various habitats, and potentially also for developing watershed-specific threshold values based on additional elements. Also, all transect had not been drawn in the maximum growth axis in all otoliths. Therefore, the distance from core, particularly the threshold value of 513 micrometer, does not always correspond to the same length and/or age of an individual (Arai et al. 2006; Denis et al. 2023). However, this approach was suitable in excluding both the Sargasso Sea birth signal and the initial coastal signal in Sr:Ca for all transects. By removing the initial 513 micrometers of data for each transect, we might filter out early habitat choices. However, we aimed at classifying active habitat choices of eel, therefore the exclusion of the initial arrival phase was needed (see also Arai et al. 2006, Teichert et al. 2023).

## Conclusion

Here, using existing otolith microchemistry data of more than 3600 European eel individuals caught along a salinity gradient from fully marine to freshwater, we show that eel display large intraspecific variation in habitat use, including coastal residents, freshwater residents and habitat-shifting life-histories, i.e. using both coastal and freshwater habitats to grow. Our findings support the growing body of scientific literature showing a high degree of individual variation in habitat utilization by European eel during its continental growth life-phase. Our findings suggest that coastal habitats are important to include when assessing the status and the population development of the European eel. Future studies should aim at assessing habitat utilization and individual variation in life-histories of European eel during pre-industrial conditions, when freshwater accessibility was not negatively affected by anthropogenic barriers to the same extent as it is today, and when survival rates in freshwater systems were less affected by hydropower turbines.

## Acknowledgements

We would like to thank the staff at the former Swedish Board of Fisheries, and at the present Swedish University of Agricultural Sciences, Department of Aquatic Recourses (SLU Aqua), that have been involved in the collection and dissection of eel over the years, leading to the important and substantial databases KUL and Sötebasen. We also thank those preparing eel otoliths and conducting otolith microchemistry analyses. In particular we want to mention Håkan Wickström and Jennie Strömquist who have been driving much of the otolith microchemistry work.

## Author contributions

Author contribution roles are listed here according to the Contributor Role Taxonomy (CRediT) guidelines (https://credit.niso.org):

Conceptualization: PJ, JS, YH

Data curation: PJ, LS, RvG, EM

Formal analysis: PJ, LS, RvG

Funding Acquisition: PJ, JS, YH

Project administration: PJ

Validation: PJ, RvG

Visualization: PJ

Writing – Original draft: PJ, LS

Writing – Review and Editing: all authors.

## Data availability

Data and R-scripts are available via Github (https://github.com/PhilipJac/FREEL_1).

## Ethical statement

Animal ethical approval was not required for this study since we analysed existing data. The original data used here had been collected between the years 1984-2024, as part of several different monitoring programs, in accordance with the animal ethical legislation valid at that time, with the latest permit being valid from 2019-2024 (Dnr. 5.8.18-10169/2019). Fisheries dependent data collection complied with fisheries regulations, and fisheries independent data collection was performed under fisheries regulation exempts.

## Supplementary Information

This supplementary information contains tables (Table S1-S3) and figures (Figure S1-S3) showing the setup-details for the laser ablation inductively coupled plasma mass spectrometry (LA-ICP-MS) at the Department of Geology at Lund University in Lund, Sweden (Table S1), the location and summary data of the eel used for determining habitat specific threshold values in Sr:Ca (Fig. S1, Table S2), and a comparison between acquired otolith microchemistry data derived using both FE-EPMA and LA-ICP-MS with overlaying transects for 25 eel individuals (Fig. S2, S3 and Table S3).

**Table S1.** Setup for the laser ablation inductively coupled plasma mass spectrometry (LA-ICP-MS) at the Department of Geology at Lund University in Lund, Sweden.

| **Laser ablation system** |  |
| --- | --- |
| Make, Model & type | Photon machines, Analyte G2 excimer laser |
| Ablation cell & volume | HelEx 2-volume cell with eQC |
| Laser wavelength | 193 nm |
| Pulse width | <4 ns |
| Fluence | 2 J/cm<sup>2</sup> on otolith and MACS-3 and 3 J/cm<sup>2</sup> on NIST glass |
| Repetition rate | 16 Hz |
| Scan speed | Varied |
| Spot size | Varied |
| Background collection | 30 seconds prior to each analysis |
| Ablation setup | Line scan; 30 seconds on standards |
| Cell carrier gas flow | 0.8 l/min He |
| **ICP-MS instrument** |  |
| Make, Model & type | Bruker Aurora Elite Quadrupole ICP-MS |
| RF power | Ca. 1300 W |
| Make-up gas flow | Ca. 0.95 l/min Ar |
| Detection system | Single collector discrete dynode electron multiplier or DDEM |
| Masses measured | <sup>7</sup>Li, <sup>11</sup>B, <sup>25</sup>Mg, <sup>26</sup>Mg, <sup>31</sup>P, <sup>43</sup>Ca, <sup>55</sup>Mn, <sup>63</sup>Cu, <sup>66</sup>Zn, <sup>88</sup>Sr, <sup>127</sup>I, <sup>137</sup>Ba, <sup>208</sup>Pb |
| **Data reduction** |  |
| Software | Iolite v3.63 |
| Primary standard | NIST SRM 610<br>MACS-3 |
| Secondary standard(s) | NIST SRM 612 |
| Internal standard | <sup>43</sup>Ca - Ca mean concentration in otolith set to 38 wt. % |

## Reference eels for determination of habitat specific Sr:Ca threshold values

To determine threshold values in Sr:Ca corresponding to growth in various salinities, we selected naturally recruited eels caught at freshwater and coastal sampling sites along the salinity gradient from Skagerrak to Kattegat to the Baltic Sea (Fig. S1, Table S2).

**Figure S1.**
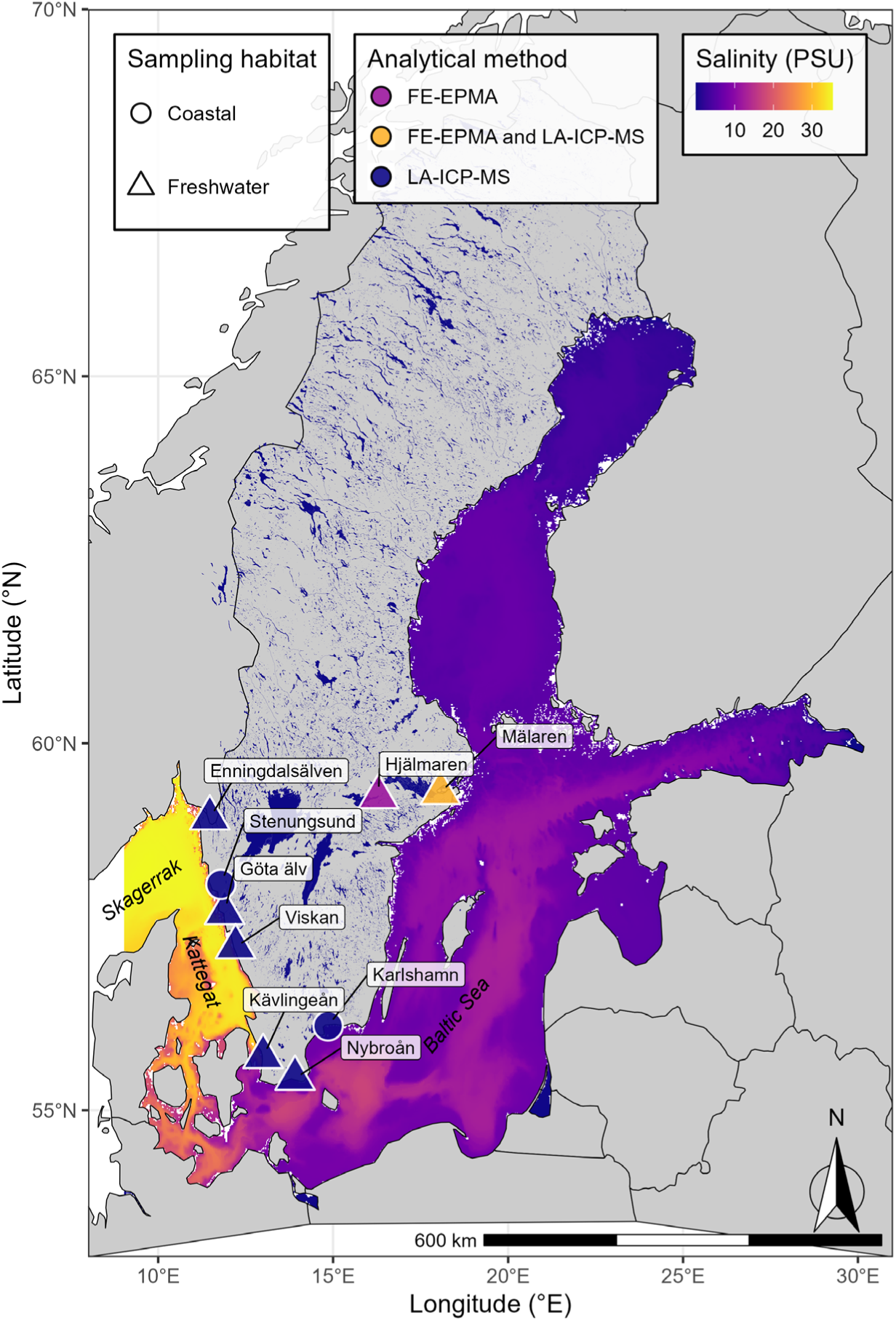
Location of the reference eel sampling sites used to derive habitat specific threshold values in Strontium to Calcium (Sr:Ca) ratios of otolith transects. Data on sea floor salinity was downloaded from the Copernicus Monitoring Environment Marine Service web portal (CMEMS 2026).

**Table S2.**
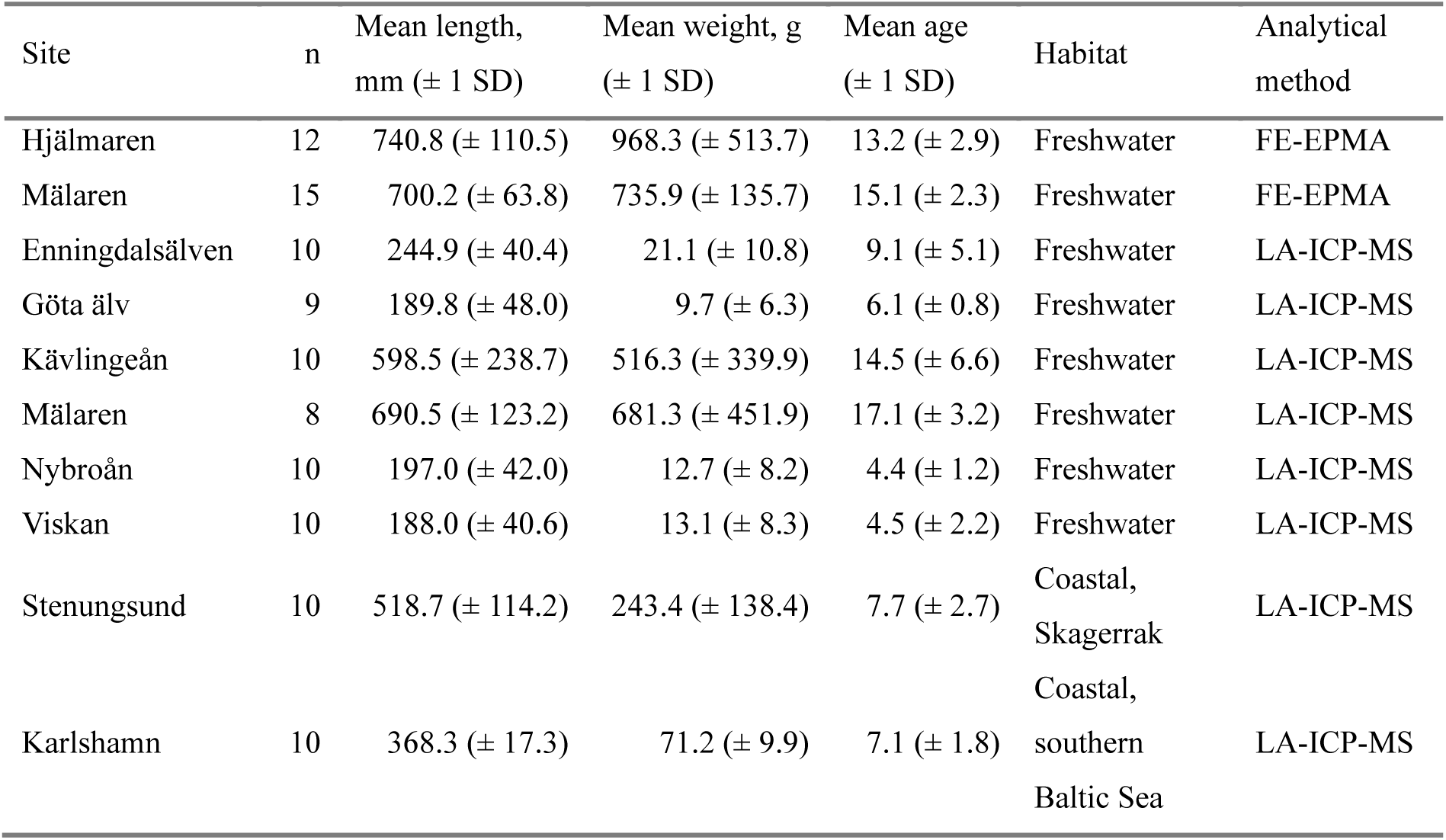
Descriptive data of the selected reference eels used to determine Sr:Ca threshold values corresponding to growth in various salinities for otolith chemistry data derived using either FE-EPMA or LA-ICP-MS otolith microchemistry data.

| Site | n | Mean length,<br>mm (± 1 SD) | Mean weight, g<br>(± 1 SD) | Mean age<br>(± 1 SD) | Habitat | Analytical<br>method |
| --- | --- | --- | --- | --- | --- | --- |
| Hjälmaren | 12 | 740.8 (± 110.5) | 968.3 (± 513.7) | 13.2 (± 2.9) | Freshwater | FE-EPMA |
| Mälaren | 15 | 700.2 (± 63.8) | 735.9 (± 135.7) | 15.1 (± 2.3) | Freshwater | FE-EPMA |
| Enningdalsälven | 10 | 244.9 (± 40.4) | 21.1 (± 10.8) | 9.1 (± 5.1) | Freshwater | LA-ICP-MS |
| Göta älv | 9 | 189.8 (± 48.0) | 9.7 (± 6.3) | 6.1 (± 0.8) | Freshwater | LA-ICP-MS |
| Kävlingeån | 10 | 598.5 (± 238.7) | 516.3 (± 339.9) | 14.5 (± 6.6) | Freshwater | LA-ICP-MS |
| Mälaren | 8 | 690.5 (± 123.2) | 681.3 (± 451.9) | 17.1 (± 3.2) | Freshwater | LA-ICP-MS |
| Nybroån | 10 | 197.0 (± 42.0) | 12.7 (± 8.2) | 4.4 (± 1.2) | Freshwater | LA-ICP-MS |
| Viskan | 10 | 188.0 (± 40.6) | 13.1 (± 8.3) | 4.5 (± 2.2) | Freshwater | LA-ICP-MS |
| Stenungsund | 10 | 518.7 (± 114.2) | 243.4 (± 138.4) | 7.7 (± 2.7) | Coastal,<br>Skagerrak | LA-ICP-MS |
| Karlshamn | 10 | 368.3 (± 17.3) | 71.2 (± 9.9) | 7.1 (± 1.8) | Coastal,<br>southern<br>Baltic Sea | LA-ICP-MS |

## Comparison between FE-EPMA and LA-ICP-MS Sr:Ca ratios for life-history assignment and quantification of habitat shifts

We compared the assigned life-history and number of habitat shifts between freshwater and coastal habitats for 25 eel individuals caught in lake Mälaren, based otolith micro chemistry data from the same otolith, acquired using both Field-Emission Electron microprobe analyses (FE-EPMA) and Laser Ablation Inductively Coupled Plasma Mass Spectrometry (LA-ICP-MS). All 25 individuals were assigned a habitat-shifting life-history independent of analytical method (Table S3), while the number of shifts between freshwater (FE-EPMA) or freshwater/estuary (LA-ICP-MS) and coastal habitats was generally higher when quantified based on LA-ICP-MS compared to FE-EPMA data (Table S3). Overall, the Sr:Ca ratio profiles were generally similar for all individuals irrespective of analytical method, all though the LA-ICP-MS profiles consists of more data-points compared to FE-EPMA transects (n=30 for FE-EPMA profiles; n= 752-1431 for LA-ICP-MS profiles) and show greater small-scale variability in the Sr:Ca ratio compared to the FE-EPMA data (Fig. S2, S3).

**Table S3.** Assigned life-history and number of shifts between freshwater/estuary and coastal habitats of 25 eel caught in lake Mälaren based on the Sr.Ca ratio derived using Field-Emission Electron microprobe analyses (FE-EPMA) and Laser Ablation Inductively Coupled Plasma Mass Spectrometry (LA-ICP-MS) analyses of the same otolith with overlaying transects.

| Individual ID-nr | Total length (mm) | Total weight (g) | Assigned life-history |  | Habitat shifts |  |
| --- | --- | --- | --- | --- | --- | --- |
|  |  |  | FE-EPMA | LA-ICP-MS | FE-EPMA | LA-ICP-MS |
| 31584 | 808 | 1343 | Shifting | Shifting | 1 | 5 |
| 31585 | 800 | 1070 | Shifting | Shifting | 1 | 15 |
| 31586 | 689 | 719 | Shifting | Shifting | 1 | 3 |
| 31587 | 776 | 945 | Shifting | Shifting | 1 | 7 |
| 31588 | 755 | 873 | Shifting | Shifting | 1 | 1 |
| 31591 | 725 | 730 | Shifting | Shifting | 1 | 3 |
| 31596 | 760 | 809 | Shifting | Shifting | 3 | 3 |
| 31599 | 831 | 1207 | Shifting | Shifting | 1 | 1 |
| 31600 | 743 | 874 | Shifting | Shifting | 3 | 3 |
| 31601 | 640 | 588 | Shifting | Shifting | 1 | 3 |
| 31602 | 695 | 687 | Shifting | Shifting | 1 | 5 |
| 31606 | 788 | 1122 | Shifting | Shifting | 3 | 5 |
| 31616 | 698 | 727 | Shifting | Shifting | 1 | 3 |
| 31617 | 789 | 948 | Shifting | Shifting | 1 | 3 |
| 31618 | 796 | 946 | Shifting | Shifting | 1 | 3 |
| 31620 | 742 | 878 | Shifting | Shifting | 1 | 5 |
| 31621 | 715 | 746 | Shifting | Shifting | 1 | 1 |
| 31623 | 700 | 740 | Shifting | Shifting | 1 | 1 |
| 31625 | 728 | 692 | Shifting | Shifting | 1 | 3 |
| 31627 | 733 | 813 | Shifting | Shifting | 1 | 9 |
| 33916 | 765 | 1012 | Shifting | Shifting | 1 | 7 |
| 33917 | 844 | 1134 | Shifting | Shifting | 1 | 7 |
| 33919 | 894 | 1464 | Shifting | Shifting | 1 | 1 |
| 33920 | 795 | 1033 | Shifting | Shifting | 1 | 7 |
| 33924 | 796 | 1057 | Shifting | Shifting | 1 | 5 |

**Figure S2.**
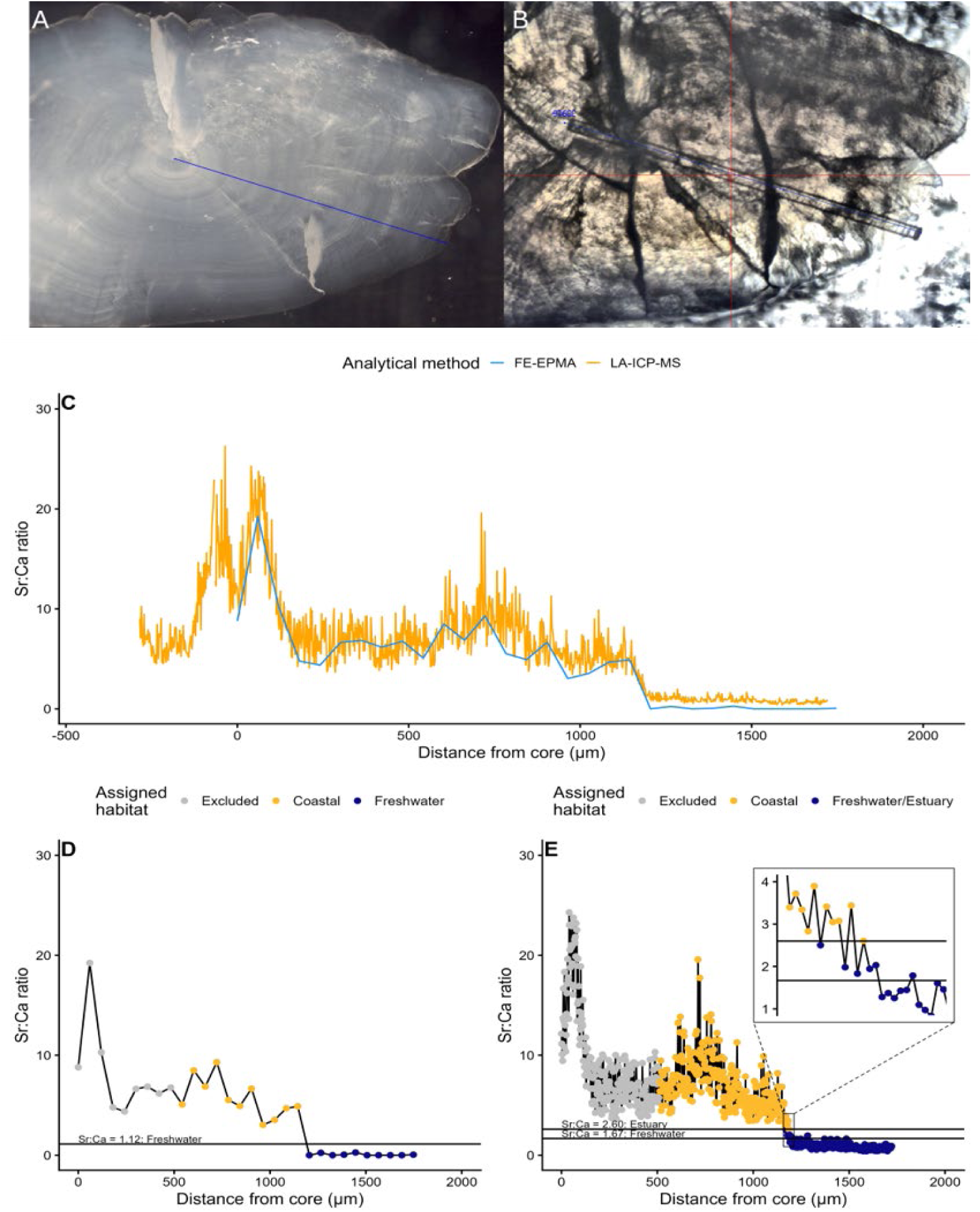
Derived Sr:Ca ratio transects of one eel otolith using either FE-EPMA (panel A, C & D) or LA-ICP-MS (panel B, C & E), with overlayed transects (Individual ID-Nr: 33916, caught in lake Mälaren). Panel A show the grinded and polished otolith in epoxy prior and the transect location on the otolith prior to the FE-EPMA analysis. Panel B show the photo of the same otolith in the LA-ICP-MS instrument and the location of the analysed transect. Panel C show the Sr:Ca ratio of the transect profiles of the FE-EPMA and the LA-ICP-MS analyses. Panel D show the assigned habitat for each data point of the transect and the horizontal FE-EPMA specific Sr:Ca ratio threshold value for freshwater habitat growth. Panel E show the assigned habitat for each data point of the transect and the horizontal LA-ICP-MS specific Sr:Ca ratio threshold value for freshwater and estuary habitat habitat growth. The inset show data-points in close proximity to the threshold values in Sr:Ca corresponding to a shift between coastal and estuary/freshwater growth.

**Figure S3.**
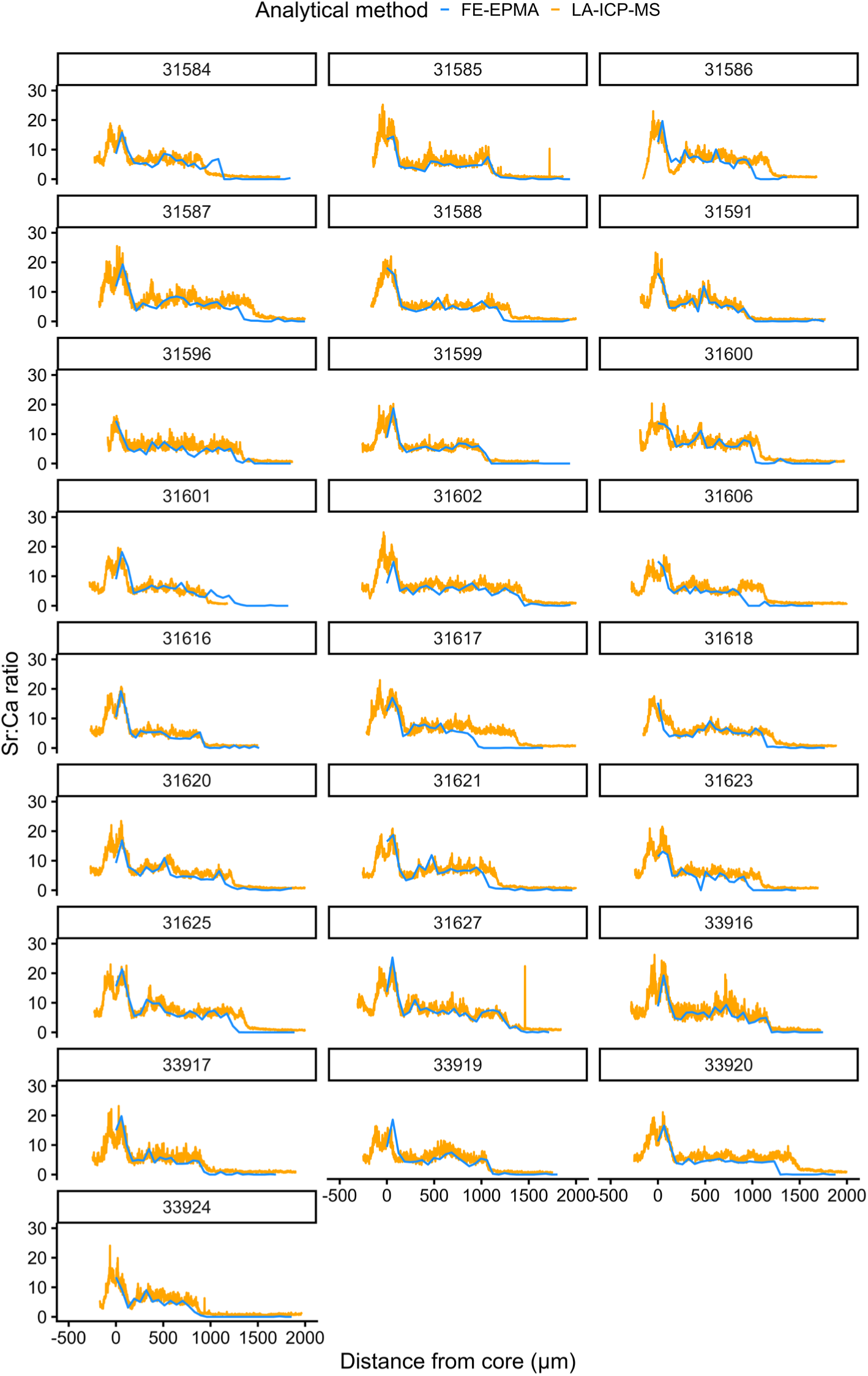
Comparison of Sr:Ca ratios of the same otolith with overlayed transects derived using FE-EPMA (blue-line, n=30) and LA-ICP-MS (orange-yellow line, n varies between 752-1431 data-points) of 25 eel caught in lake Mälaren where each panel show data for each individual and its corresponding Individual ID nr.

## References

1. Arai T, Kotake A, McCarthy TK. 2006. Habitat use by the European eel *Anguilla anguilla* in Irish waters. Estuarine, Coastal and Shelf Science, 67, 4: 569–578. DOI: 10.1016/j.ecss.2006.01.001.

2. Berg, T., Steen, JB. 1965. Physiological mechanisms for aerial respiration in the eel. Comparative Biochemistry and Physiology, 15, 4: 469–484. DOI: 10.1016/0010-406X(65)90147-7.

3. Bonsdorff, E. 2006. Zoobenthic diversity-gradients in the Baltic Sea: Continuous post-glacial succession in a stressed ecosystem. Journal of Experimental Marine Biology and Ecology, 330, 1: 383–391. DOI: 10.1016/j.jembe.2005.12.041.

4. Calles, O., Olsson, I.C., Comoglio, C., Kemp, P.S., Blunden, L., Schmitz, M. and Greenberg, L.A. 2010. Size-dependent mortality of migratory silver eels at a hydropower plant, and implications for escapement to the sea. Freshwater Biology, 55: 2167–2180. DOI: 10.1111/j.1365-2427.2010.02459.x

5. Campana, S. E. 1999. Chemistry and composition of fish otoliths: pathways, mechanisms and applications. Marine Ecology Progress Series, 188: 263–297. DOI: 10.3354/meps188263

6. Campana, S. E., & Thorrold, S. R. 2001. Otoliths, increments, and elements: keys to a comprehensive understanding of fish populations? Canadian Journal of Fisheries and Aquatic Sciences, 58, 1: 30–38. DOI: 10.1139/f00-177

7. Campitelli, E. 2025. ggnewscale: Multiple fill and colour scales in ‘ggplot2. URL: https://CRAN.R-project.org/package=ggnewscale

8. Capoccioni, F., Lin, DY., Iizuka, Y., Tzeng, WN., Ciccotti, E. 2013. Phenotypic plasticity in habitat use and growth of the European eel (*Anguilla anguilla*) in transitional waters in the Mediterranean area. Ecology of Freshwater Fish. 23, 1: 65–76. DOI: 10.1111/eff.12049

9. Chang, W. 2023. webshot2: Take Screenshots of Web Pages. URL: https://CRAN.R-project.org/package=webshot2

10. Clevestam, P. D., & Wickström, H. 2008. Rädda ålen och ålfisket! – Ett nationellt bidrag till en europeisk förvaltningsplan. Swedish Board of Fisheries, Drottningholm. 1–42. [in Swedish]

11. CMEMS 2026. Copernicus Monitoring Environment Marine Service. Accessed 2026-03-15. URL: https://data.marine.copernicus.eu/product/BALTICSEA_MULTIYEAR_PHY_003_011/description, DOI: 10.48670/moi-00013.

12. Dadswell, M., Spares, A., Reader, J., McLean, M., McDermott, T., Samways, K., & Lilly, J. 2022. The decline and impending collapse of the Atlantic Salmon (*Salmo salar*) population in the North Atlantic Ocean: A review of possible causes. Reviews in Fisheries Science & Aquaculture, 30, 2: 215–258. DOI: 10.1080/23308249.2021.1937044

13. Darwall, W., Bremerich, V., De Wever, A., et al. 2018. The alliance for freshwater life: A global call to unite efforts for freshwater biodiversity science and conservation. Aquatic Conservation: Marine and Freshwater Ecosystems 28: 1015–1022. DOI: 10.1002/aqc.2958

14. Daverat, F., Limburg, K. E., Thibault, I., Shiao J., Dodson, J.J., Caron, F., Tzeng, W.N., Iizuka, Y., Wickström, H. 2006. Phenotypic plasticity of habitat use by three temperate eel species, Anguilla anguilla, A. japonica and A. rostrata. Marine Ecology Progress Series 308: 231–241. DOI: 10.3354/meps308231

15. Daverat, F., Tomás, J. 2006. Tactics and demographic attributes in the European eel *Anguilla anguilla* in the Gironde watershed, SW France. Marine Ecology Progress Series. 307: 247–257. DOI: 10.3354/meps307247

16. Dempson, J. B., Van Leeuwen, T. E., Bradbury, I. R., Lehnert, S. J., Coté, D., Cyr, F., Pretty, C., Kelly, N. I. 2024. A review of factors potentially contributing to the long-term decline of Atlantic Salmon in the Conne River, Newfoundland, Canada. Reviews in Fisheries Science & Aquaculture, 32, 3: 479–504. DOI: 10.1080/23308249.2024.2341023

17. Denis, J., Mahé, K., Tabouret, H., Rabhi, K., Boutin, K., Diop, M., Amara, R. 2023. Relationship between habitat use and individual condition of European eel (*Anguilla anguilla*) in six estuaries of the eastern English Channel (North-eastern Atlantic ocean). Estuarine, Coastal and Shelf Science, 291: 108446. DOI: 10.1016/j.ecss.2023.108446

18. Dodson, J.J., Aubin-Horth, N., Thériault, V., Páez, D.J. 2013. The evolutionary ecology of alternative migratory tactics in salmonid fishes. Biological Reviews, 88, 3: 602–615. DOI: 10.1111/brv.12019

19. Duarte, G., Segurado, P., Haidvogl, G., Pont, D., Ferreira, M.T., Branco, P. 2021. Damn those damn dams: Fluvial longitudinal connectivity impairment for European diadromous fish throughout the 20th century. Science of The Total Environment 761: 143293. DOI: 10.1016/j.scitotenv.2020.143293.

20. Dunnington, D. 2023. ggspatial: Spatial data framework for ggplot2. URL: https://CRAN.R-project.org/package=ggspatial

21. Durif, C., Arts, M., Bertolini, F., Cresci, A., Daverat, F., Karlsbakk, E., Koprivnikar, J., Moland, E., Olsen, E.M., Parzanini, C., Power, M., Rohtla, M., Skiftesvik, A.B., Thorstad, E., Vøllestad, L.A., Browman, H. 2023. The evolving story of catadromy in the European eel (*Anguilla anguilla*). ICES Journal of Marine Science, 80, 9: 2253–2265. DOI: 10.1093/icesjms/fsad149

22. Édeline, E., Élie, P., 2004. Is salinity choice related to growth in juvenile eel *Anguilla anguilla*? Cybium, International Journal of Ichtyology, 28:77–82.

23. Enbody, E. D., Pettersson, M. E., Sprehn, C. G., Palm, S., Wickström, H., Andersson, L. 2021. Ecological adaptation in European eels is based on phenotypic plasticity. Proceedings of the National Academy of Sciences, 118, 4: e2022620118. DOI: 10.1073/pnas.2022620118

24. Fabosa 2002. Fish ageing by otolith shape analysis: Final report to the European Commission (EC contract no. FAIR CT97 3402): 1–204. URL: https://www.ices.dk/community/documents/pgccdbs/fabosa_rapp02_hel.pdf

25. Garnier, S., Ross, N., Rudis, R., Camargo, A.P., Sciaini, M., Scherer, C. 2024. viridis(Lite) – Colorblind-Friendly Color Maps for R. DOI: 10.5281/zenodo.4679423

26. Gillson, J.P., Bašić, T., Davison, P.I. et al. 2022. A review of marine stressors impacting Atlantic salmon *Salmo salar*, with an assessment of the major threats to English stocks. Reviews in Fish Biology & Fisheries, 32: 879–919. DOI: 10.1007/s11160-022-09714-x

27. Gohel, D., Moog, S. 2024. officer: Manipulation of Microsoft Word and PowerPoint Documents URL: https://CRAN.R-project.org/package=officer.

28. Gohel, D., Skintzos, P. 2024. flextable: Functions for Tabular Reporting. URL: https://CRAN.R-project.org/package=flextable.

29. Gross, M.R., Coleman, R.M., McDowall, R.M. 1988. Aquatic productivity and the evolution of diadromous fish migration. Science, 239: 1291–1293. DOI: 10.1126/science.239.4845.1291

30. Guiry, E., Robson, H.K. 2024. Deep antiquity of seagrasses supporting European eel fisheries in the western Baltic. Proceedings of the Royal Society B: Biological Sciences, 291: 20240674. DOI: 10.1098/rspb.2024.0674

31. Hamer, P., Henderson, A., Hutchison, M., Kemp, J., Green, C., Feutry, P. 2015. Atypical correlation of otolith strontium:calcium and barium:calcium across a marine–freshwater life history transition of a diadromous fish. Marine & Freshwater Research 66: 411–419. DOI: 10.1071/MF14001

32. Hellström, J., Paton, C., Woodhead, J., Hergt, J. 2008. Iolite: Software for spatially resolved LA-(quad and MC)-ICP-MS analysis. In P. Sylvester (Ed.), Laser ablation ICP–MS in the earth sciences: Current practices and outstanding issues: 343–348. Mineralogical Association of Canada Short Course Series.

33. Hester J, Wickham H, Gjoneski O. 2024. odbc: Connect to ODBC Compatible Databases (using the DBI Interface). URL: https://CRAN.R-project.org/package=odbc.

34. Hijmans, R. 2023. terra: Spatial data analysis. URL: https://CRAN.R-project.org/package=terra.

35. Horton, T.W., Block, B.A., Drumm, A., Hawkes, L.A, O’Cuaig, M., Ó Maoiléidigh, N., O’Neill, R., Schallert, R.J., Stokesbury, M.J.W., Witt, M.J. 2020. Tracking Atlantic bluefin tuna from foraging grounds off the west coast of Ireland. ICES Journal of Marine Science, 77, 6: 2066–2077. DOI: 10.1093/icesjms/fsaa090

36. Hugh-Jones, D. 2024. ggmagnify: Create a magnified inset of part of a ggplot object. URL: https://github.com/hughjonesd/ggmagnify.

37. Ibbotson, A., Smith, J., Scarlett, P., Aprhamian, M. 2002. Colonisation of freshwater habitats by the European eel *Anguilla anguilla*. Freshwater Biology, 47: 1696–1706. DOI: 10.1046/j.1365-2427.2002.00930.x

38. ICES 2025. Joint EIFAAC/ICES/GFCM Working Group on Eels (WGEEL). ICES Scientific Reports, 7, 99: 1–134. DOI: 10.17895/ices.pub.30488120.v2

39. Jacobson, P., Gårdmark, A., Huss, M. 2019. Population and size-specific distribution of Atlantic salmon *Salmo salar* in the Baltic Sea over five decades. Journal of Fish Biology, 96, 2: 408–417. DOI: 10.1111/jfb.14213

40. Kettle, A.J., Heinrich, D., Barrett, J.H., Benecke, N., Locker, A. 2008. Past distributions of the European freshwater eel from archaeological and palaeontological evidence. Quaternary Science Reviews, 27, 13–14: 1309–1334. DOI: 10.1016/j.quascirev.2008.03.005.

41. Klemetsen, A., Amundsen, P.A., Dempson, J.B., Jonsson, B., Jonsson, N., O’Connell, M.F., Mortensen, E. 2003. Atlantic salmon Salmo salar L., brown trout Salmo trutta L. and Arctic charr Salvelinus alpinus (L.): a review of aspects of their life histories. Ecology of Freshwater Fish, 12, 1: 1–59. DOI: 10.1034/j.1600-0633.2003.00010.x

42. Klemetsen, A. 2013. The most variable vertebrate on Earth. Journal of Ichthyol. 53: 781–791. DOI: 10.1134/S0032945213100044

43. Knight, T., McCoy, M., Chase, J. et al. 2005. Trophic cascades across ecosystems. Nature 437: 880–883. DOI: 10.1038/nature03962

44. KUL 2026. The database for fish monitoring fish along the coast – KUL. Swedish University of Agricultural Sciences (SLU), Department of Aquatic Resources. https://slu.se/kul. [access date 2026-02-12]

45. Limburg, K. E., Wickström, H., Svedäng, H., Elfman, M., & Kristiansson, P. 2003. Do stocked freshwater eels migrate? Evidence from the Baltic suggests “Yes”. American Fisheries Society Symposium, 33: 275–284.

46. Limburg, K. E., Waldman, J. R. 2009. Dramatic declines in North Atlantic diadromous fishes. BioScience, 59, 11: 955–965. DOI: 10.1525/bio.2009.59.11.7

47. Linton, E. D., Jónsson, B., Noakes, D. L. 2007. Effects of water temperature on the swimming and climbing behaviour of glass eels, *Anguilla* spp. Environmental Biology of Fishes, 78, 3: 189–192. DOI: 10.1007/s10641-005-1367-9

48. Magnusson, A. K., & Dekker, W. 2021. Economic development in times of population decline – A century of European eel fishing on the Swedish west coast. ICES Journal of Marine Science, 78, 1: 185–198. DOI: 10.1093/icesjms/fsaa213

49. Marohn, L., Hilge V, Zumholz K, Klügel A, Anders H, Hanel R. 2011. Temperature dependency of element incorporation into European eel (*Anguilla anguilla*) otoliths. Analytical and Bioanalytical Chemistry 399: 2175–2184. DOI: 10.1007/s00216-010-4412-2

50. Massicotte, P., South, A. 2023. rnaturalearth: World map data from natural earth. URL: https://CRAN.R-project.org/package=rnaturalearth

51. McDowall, R.M. 1999. Different kinds of diadromy: Different kinds of conservation problems. ICES Journal of Marine Science, 56, 4: 410–413. DOI: 10.1006/jmsc.1999.0450

52. Miller, T.E., Rudolf, V.H. 2011. Thinking inside the box: community-level consequences of stage-structured populations. Trends in ecology & evolution, 26, 9: 457–466. DOI: 10.1016/j.tree.2011.05.005

53. Miller, M.J., Westerberg, H., Sparholt, H., Wysujack, K., Sørensen, S.R., Marohn, L., Jacobsen, M.W, Freese, M., Ayala, D.J., Pohlmann, J.D., Svendsen, J.C., Watanabe, S., Andersen, L., Møller, P.R., Tsukamoto, K., Munk, P., Hanel, R. 2019. Spawning by the European eel across 2000 km of the Sargasso Sea. Biology Letters, 15, 4: 20180835. DOI: 10.1098/rsbl.2018.0835

54. Moore, J.W., Yeakel, J.D., Peard, D., Lough, J., Beere, M. 2014. Life-history diversity and its importance to population stability and persistence of a migratory fish: steelhead in two large North American watersheds. Journal of Animal Ecology, 83, 5: 1035–1046. DOI: 10.1111/1365-2656.12212

55. Moriarty, C., Dekker, W. 1997. Management of the European Eel. Second report of EC Concerted Action AIR A94–1939.

56. Nilsson, C., Reidy, C.A., Dynesius, M., Revenga, C. 2005. Fragmentation and flow regulation of the world’s large river systems. Science, 308: 405–408. DOI: 10.1126/science.1107887.

57. Ohlberger, J., Buhle, E.R., Buehrens, T.W., Kendall, N.W., Harbison, T., Claiborne, A.M., Losee, J.P., Whitney, J., Scheuerell, M.D. 2025. Declining marine survival of Steelhead trout linked to climate and ecosystem change. Fish and Fisheries, 26, 3: 331–345. DOI: 10.1111/faf.12878

58. OpenAI. 2026. ChatGPT (May 2026 free version) [Large language model]. https://openai.com/

59. Palm, S., Dannewitz, J., Prestegaard, T., & Wickström, H. 2009. Panmixia in European eel revisited: no genetic difference between maturing adults from southern and northern Europe. Heredity 103: 82–89. DOI: 10.1038/hdy.2009.51

60. Patin, R., Etienne, P. M, Lebarbier, E., Chamaille-Jammes, S., Benhamou &, S. 2020. Identifying stationary phases in multivariate time series for highlighting behavioural modes and home range settlements. Journal of Animal Ecology, 89,1: 44–56. DOI: 10.1111/1365-2656.13105

61. Paton, C., Hellström, J., Paul, B., Woodhead, J., & Hergt, J. 2011. Iolite: Freeware for the visualisation and processing of mass spectrometric data. Journal of Analytical Atomic Spectrometry, 26, 12: 2508–2518. DOI: 10.1039/C1JA10172B

62. Pebesma, E., & Bivand, R. 2023. Spatial Data Science: With Applications in R. Chapman and Hall/CRC. DOI: 10.1201/9780429459016

63. Pebesma, E., 2018. Simple Features for R: Standardized Support for Spatial Vector Data. The R Journal 10, 1: 439–446. DOI: 10.32614/RJ-2018-009

64. Pedersen, T. 2024. patchwork: The composer of plots. URL: https://CRAN.R-project.org/package=patchwork.

65. Persson, L., DeRoos, A.M. 2003. Adaptive habitat use in size-structures populations: Linking individual behaviour to population Processes. Ecology, 84, 5: 1129–1139. DOI: 10.1890/0012-9658(2003)084[1129:AHUISP]2.0.CO;2

66. Pike, C., Crook, V., Gollock, M. 2020. Anguilla anguilla. The IUCN Red List of Threatened Species 2020: e.T60344A152845. DOI: 10.2305/IUCN.UK.2020-2.RLTS.T60344A152845178.en

67. Poff, N.L., Olden, J.D., Merritt, D.M., Pepin, D.M. 2007. Homogenization of regional river dynamics by dams and global biodiversity implications. Proceedings of the National Academy of Sciences, 104, 14: 5732–5737. DOI: 10.1073/pnas.0609812104

68. Quinn, T.P., Myers, K.W. 2004. Anadromy and the marine migrations of Pacific salmon and trout: Rounsefell revisited. Reviews in Fish Biology and Fisheries, 14: 421–442. DOI: 10.1007/s11160-005-0802-5

69. R Core Team. 2025. R: A language and environment for statistical computing [Computer software]. R Foundation for Statistical Computing. https://www.R-project.org/

70. R Special Interest Group on Databases, Wickham H, Müller K. 2024. DBI: R Database Interface, URL: https://CRAN.R-project.org/package=DBI.

71. Righton, D., Westerberg, H., Feunteun, E., Økland, F., Gargan, P., Amilhat, E., Metcalfe, J., Lobon-Cervia, J., Sjöberg, N., Simon, J., Acou, A., Vedor, M., Walker, A., Trancart, T., Brämick, U., & Aarestrup, K. 2016. Empirical observations of the spawning migration of European eels: The long and dangerous road to the Sargasso Sea. Science Advances, 2, 10: e1501694. DOI: 10.1126/sciadv.1501694

72. Rikardsen, A.H., Righton, D., Strøm, J.F. et al. 2021. Redefining the oceanic distribution of Atlantic salmon. Scientific Reports 11, 1: 12266. DOI: 10.1038/s41598-021-91137-y

73. Rothla, M., Daverat, F., Arts, M.T., Browman, H.I., Parzanini, C., Skiftesvik, A.B., Thorstad, E.B., van der Meeren, T., Vøllestad, L.A., Durif, C. 2023. Habitat use and growth of yellow-stage European eel in coastal and freshwater ecosystems in Norway. Canadian Journal of Fisheries and Aquatic Sciences, 80, 1: 14–26. DOI: 10.1139/cjfas-2022-0033

74. Schmidt, J. 1912. The reproduction and spawning places of the freshwater eel (*Anguilla vulgaris*). Nature 89: 633–636. DOI: 10.1038/089633a0

75. Schmidt, J. 1922. The Breeding Places of the Eel. Philosophical Transactions of the Royal Society of London, Series B 211: 179–208. DOI: 10.1098/rstb.1923.0004

76. Schreiber, S., Rudolf, V.H.W. 2008. Crossing habitat boundaries: coupling dynamics of ecosystems through complex life cycles. Ecology Letters, 11, 6: 576–587. DOI: 10.1111/j.1461-0248.2008.01171.x

77. Silge, J., Robinson, D. 2016. tidytext: Text mining and analysis using tidy data principles in R. The Journal of Open Source Software, 1, 3: 37. DOI: 10.21105/joss.00037

78. Slowikowski, K. 2023. ggrepel: Automatically position non-overlapping text labels with ‘ggplot2’. URL: https://CRAN.R-project.org/package=ggrepel

79. SMHI. 2025. Swedish Meteorological and Hydrological Institute. URL: https://www.smhi.se/kunskapsbanken/hydrologi/hydrografiska-data/sveriges-huvudavrinningsomraden

80. Säterberg, T., Gilljam, D., Holliland, P. B., Jacobson, P., van Gemert, R. 2025. Effects of a fishery closure on the European eel stock on the Swedish west coast. Fisheries Research, 291: 107564. DOI: 10.1016/j.fishres.2025.107564

81. Tahri, M., & Panfili, J. 2023. 13-year population survey of the critically endangered European eel in the southern Mediterranean region (Algeria). Journal of Fish Biology, 102, 6: 1492–1502. DOI: 10.1111/jfb.15396

82. Tamario, C., Calles, O., Watz, J., Nilsson, P.A., Degerman, E. 2019. Coastal river connectivity and the distribution of ascending juvenile European eel *(Anguilla anguilla* L.): Implications for conservation strategies regarding fish-passage solutions. Aquatic Conservation: Marine & Freshwater Ecosystems, 29: 612–622. DOI: 10.1002/aqc.3064

83. Teichert, N., Bourillon, B., Suzuki, K. et al. 2023. Biogeographical snapshot of life-history traits of European silver eels: insights from otolith microchemistry. Aquatic Science 85, 2: 39. DOI: 10.1007/s00027-023-00940-4

84. Tesch, F. W. 2003. The Eel (R. J. White, Trans.; J. E. Thorpe, Ed.) (5th ed.). Wiley-Blackwell.

85. Thorstad, E.B., Bliss, D., Breau, C., Damon-Randall, K., Sundt-Hansen, L.E., Hatfield, E.M.C., Horsburgh, G., Hansen, H. Ó., Maoiléidigh, N., Sheehan, T., Sutton, S.G. 2021. Atlantic salmon in a rapidly changing environment – Facing the challenges of reduced marine survival and climate change. Aquatic Conservation: Marine and Freshwater Ecosystems, 31, 9: 2654–2665. DOI: 10.1002/aqc.3624

86. Tzeng, W.N., Severin, K.P., Wickström, H. 1997. Use of otolith microchemistry to investigate the environmental history of European eel *Anguilla anguilla*. Marine Ecology Progress Series, 149: 73–81. DOI: 10.3354/meps149073

87. Waldman, J.R., Quinn, T.P. 2022 North American diadromous fishes: Drivers of decline and potential for recovery in the Anthropocene. Scientific Advances, 8: eabl5486. DOI: 10.1126/sciadv.abl5486

88. Walther, B. D., & Limburg, K. E. 2012. The use of otolith chemistry to characterize diadromous migrations. Journal of Fish Biology, 81, 2: 796–825. DOI: 10.1111/j.1095-8649.2012.03371.x

89. Werner, E. E., & Gilliam, J. F. 1984. The ontogenetic niche and species interactions in size-structured populations. Annual Review of Ecology and Systematics, 15: 393–425. DOI: http://www.jstor.org/stable/2096954

90. Wickham H, Averick M, Bryan J, Chang W, McGowan LD, François R, Grolemund G, Hayes A, Henry L, Hester J, Kuhn M, Pedersen TL, Miller E, Bache SM, Müller K, Ooms J, Robinson D, Seidel DP, Spinu V, Takahashi K, Vaughan D, Wilke C, Woo K, Yutani H. 2019. Welcome to the tidyverse. Journal of Open Source Software, 4, 43: 1686. DOI: 10.21105/joss.01686

91. Wickström, H., Sjöberg, N. B. 2014. Traceability of stocked eels–the Swedish approach. Ecology of Freshwater Fish, 23, 1: 33–39. DOI: 10.1111/eff.12053

92. Wilke, C, & Wiernik, B. 2022. ggtext: Improved text rendering support for ggplot2. URL: https://CRAN.R-project.org/package=ggtext.

93. Wright, R.M., Piper, A.T., Aarestrup, K. et al. 2022. First direct evidence of adult European eels migrating to their breeding place in the Sargasso Sea. Scientific Report 12: 15362. DOI: 10.1038/s41598-022-19248-8

94. Zeileis, A., Grothendieck, G. 2005. zoo: S3 Infrastructure for regular and irregular time series. Journal of Statistical Software, 14, 6: 1–27. DOI: 10.18637/jss.v014.i06

